# KREH2 helicase represses ND7 mRNA editing in procyclic-stage *Trypanosoma brucei* by opposite modulation of canonical and “moonlighting” gRNA utilization creating a proposed mRNA structure

**DOI:** 10.1101/2024.05.06.592425

**Authors:** Joshua Meehan, Alasdair Ivens, Scott Grote, Tyler Rodshagen, Zihao Chen, Cody Goode, Sunil K. Sharma, Vikas Kumar, Addison Frese, Zachary Goodall, Laura McCleskey, Rebecca Sechrist, Lanying Zeng, Nicholas J. Savill, Silvi Rouskin, Achim Schnaufer, Suzanne M. McDermott, Jorge Cruz-Reyes

## Abstract

Unknown factors regulate mitochondrial U-insertion/deletion (U-indel) RNA editing in procyclic-form (PCF) and bloodstream-form (BSF) *T. brucei*. This editing, directed by anti-sense gRNAs, creates canonical protein-encoding mRNAs and may developmentally control respiration. Canonical editing by gRNAs that specify protein-encoding mRNA sequences occurs amid massive non-canonical editing of unclear sources and biological significance. We found PCF-specific repression at a major early checkpoint in mRNA ND7, involving helicase KREH2-dependent opposite modulation of canonical and non-canonical “terminator” gRNA utilization. Terminator-programmed editing derails canonical editing and installs proposed repressive structure in 30% of the ND7 transcriptome. BSF-to-PCF differentiation *in vitro* recreated this negative control. Remarkably, KREH2-RNAi knockdown relieved repression and increased editing progression by reverting canonical/terminator gRNA utilization. ND7 transcripts lacking early terminator-directed editing in PCF exhibited similar negative editing control along the mRNA sequence, suggesting global modulation of gRNA utilization fidelity. The terminator is a “moonlighting” gRNA also associated with mRNA COX3 canonical editing, so the gRNA transcriptome seems multifunctional. Thus, KREH2 is the first identified repressor in developmental editing control. This and our prior work support a model whereby KREH2 activates or represses editing in a stage and substrate-specific manner. KREH2’s novel dual role tunes mitochondrial gene expression in either direction during development.

## INTRODUCTION

*Trypanosoma brucei,* a member of the protist group Euglenozoa, causes Human African Trypanosomiasis (HAT) (1–3). Its complex life cycle includes procyclic forms (PCF) and bloodstream forms (BSF), which infect insect and mammalian hosts, respectively. Trypanosomatids, including *T. brucei*, exhibit massive post-transcriptional site-specific uridine insertion and deletion (U-indel) RNA editing in their large mitochondrion. Twelve of the 18 primary mRNA transcripts in mitochondria (mt) lack the correct open reading frame (ORF), which is created via U-indel RNA editing directed by trans-acting cognate guide RNAs (gRNAs), except for one case where a gRNA is provided in cis (4,5). The editing process is developmentally regulated; however, the key regulatory factors have not been defined. Pre-mRNA and gRNA transcripts are encoded in the mt-genomes of trypanosomatids (aka kinetoplast, kDNA), a planar network of catenated relaxed circles (6,7). Maxicircles encode rRNAs, and pre-mRNAs for ribosomal protein S12 (RPS12), ATPase subunit 6 (A6), and proteins in respiratory complexes. Minicircles encode hundreds of cognate gRNAs (∼45-60nt) that partially hybridize with pre-mRNA but fully complement the canonical edited sequence of mature mRNAs. A short anchor region in gRNAs initiates binding with mRNA through Watson-Crick pairing, while a guiding region directs the U-indels via Watson-Crick and GU wobble pairing (8,9). Often gRNAs contain encoded 5’ and 3’ terminal bases not used in canonical editing and a post-transcriptionally added 3’ oligo(U) tail. Editing progresses 3’ to 5’ in overlapping blocks, and each block is directed by a specific gRNA; however, redundancy is typical with multiple gRNA species covering the same block (8,10). RT-PCR of the 5’-most block typically examines the accumulation of fully edited mRNA, while detailed editing progression studies, including at early 3’ blocks, require nucleotide-resolution RNA-seq (11–13). Most mt-mRNAs are edited extensively, while others are edited moderately or are never edited (14). However, only a few mRNA molecules match the canonical pattern at steady-state, while the vast majority carry unexpected non-canonical edits of unclear function or origin. Non-canonical U-indels are usually found in editing junctions between 3’ canonical and 5’ pre-edited sequences (15,16) (**Supplementary Table ST1**, glossary of terms).

*T. brucei* uses dramatically different strategies in ATP production during development (17–21). PCF cells employ cytochrome-mediated oxidative phosphorylation. However, BSF cells employ glycolysis as they lack cytochromes and some Krebs cycle enzymes, and sugar is plentiful in serum. Accordingly, cytochrome-encoding mRNAs (complexes III and IV) with the canonical edited sequence accumulate in PCF but are rarely detectable in BSF. Other edited mRNAs, e.g., for complex I (NADH dehydrogenase), exhibit large differences in abundance between the two stages. Some transcripts, e.g., RPS12 or A6 in the F1FO-ATPase complex (complex V), are efficiently edited during all stages of development. This switch in bioenergetics may involve the regulation of editing rather than substrate availability since pre-mRNA and gRNA transcripts are relatively constant at steady-state (5,10,22).

Most of the about 40 known proteins in the editosome holoenzyme are arranged in a few ribonucleoprotein (RNP) complexes and their variants (4,15). Three <u>R</u>NA <u>E</u>diting <u>C</u>atalytic <u>C</u>omplexes (RECCs) catalyze endonucleolytic cleavage, U indels, and ligation (23–26). <u>R</u>NA <u>E</u>diting <u>S</u>ubstrate Binding <u>C</u>omplexes (RESCs) (27–30) serve as platforms for mRNA-gRNA hybrid formation and processing (31–35). We and others initially showed that purified RESC is enriched with editing substrates and products (31,33). Two RESC cryoEM reconstructed structures (RESC-A and RESC-B) may represent gRNA storage and mRNA-gRNA duplex substrate complexes, respectively (36,37). <u>R</u>NA <u>E</u>diting Helicase KRE<u>H2</u>-associated <u>C</u>omplex (REH2C) (4,5,15) contains three core proteins: the typifying DEAH-Box RNA helicase KREH2 and two directly bound helicase factors, the zinc finger protein KH2F1 and KH2F2. KREH2 controls editing fidelity (i.e., the ratio of non-canonical /canonical edits), including the first example of differential gRNA-directed non-canonical editing in PCF and BSF (38) (**Supplementary Table ST1**, glossary of terms). Sedimentation analyses suggested that REH2C may also have different variants (39). Multiple other factors that are not stably associated with the major editing complexes are also required for complete editing. These include a second RNA editing helicase, DEAD-box KREH1, which appears to promote initiator gRNA utilization (40).

Several studies indicate that REH2C is an RNP, and its association with other mt factors could involve KREH2-dependent holo-editosome remodeling. Isolated KREH2 co-purifies with proteins in RESC and RECC, KREH1, and other mt factors via RNA (31,32,41,42). All three core proteins in REH2C co-purified with mRNA substrates and products (fully and partially edited) in the absence of RESC, i.e., upon depletion of RESC1 and gRNA (32). Conversely, several purifications of RESC proteins have detected KREH2 (27–30), supporting the speculation that KREH2 could mediate remodeling between RESC-A and RESC-B CryoEM reconstructions (36). KREH2 immunoprecipitated from mt-extract or recombinantly expressed, crosslinked with RNA, and supported dsRNA unwinding *in vitro* (41,42). Inactivating point mutations in the KREH2 helicase core and dsRNA binding (dsRBD) motifs dissociated KREH2 from RESC, suggesting that a functional helicase mediates REH2C-RESC interaction (32,41,42). Furthermore, KREH2 RNAi-knockdown decreased the accumulation of canonically edited mRNAs examined by RT-qPCR in PCF and BSF (13,32,39,42), decreased editing fidelity in A6 and RPS12 in mtRNA or bound to RESC in deep sequencing studies (13,38), and shifted RESC proteins in a sedimentation analysis (43). A developmental study revealed PCF-specific KREH2 upregulation of a noncognate gRNA usage, which introduces a non-canonical 3’ High-Frequency Element (3’ HFE) in >30% of the A6 transcriptome. This 3’ HFE blocked all canonical editing in the targeted molecules, including by the initiator gRNA in A6 (38). The latter observation prompted us to test the hypothesis that KREH2 modulates the editing of substrates that are known to be developmentally regulated.

In this study, we examined mRNA ND7 (NADH dehydrogenase subunit 7), which includes two editing domains. The ND7 3’-domain (ND7 3’) exhibits preferential maturation in BSF cells by unknown mechanisms (14,44,45), including upon induced *in vitro* differentiation from PCF into mammalian infective forms (46); however, differences using an RT-PCR assay were also reported between strains (46,47). Here, nucleotide-resolution mtRNA-seq in our lab strains revealed that KREH2 represses ND7 at an early editing checkpoint. Namely, KREH2 conversely inhibited cognate gRNA and promoted non-cognate “terminator” gRNA utilization at this checkpoint. Efficient repression in native RESC6-purified RESC complexes suggests the involvement of active editosomes. This novel terminator installs a structural 3’ HFE in ∼30% of the ND7 transcriptome that hinders canonical editing progression. Notably, KREH2 loss-of-function increased editing fidelity and progression. An increase in ND7 canonical editing upon KREH2 loss is unprecedented and challenges the original definition of an editing factor. Loss of KREH2 also induced similar converse changes between canonical and non-canonical edits along the length of ND7 transcripts that did not receive the 3’ HFE in PCF cells. Thus, KREH2 is the first known factor able to repress editing and regulate editing fidelity by governing global gRNA utilization. Our current studies on ND7 and prior work in other mRNAs (13,38) support a general regulatory model in which KREH2 has an unprecedented dual role able to stimulate or repress editing. In this duality, KREH2 controls the function of gRNAs, which specify both canonical or novel regulatory editing, in a substrate- and stage-dependent manner to modulate gene expression in any direction during development.

## MATERIAL AND METHODS

### General PCF and BSF cell culture and transfection

*T. brucei* cell lines in this study, strains Lister 427 29-13 PCF and Lister 427 Single Marker BSF (monomorphic), each expressing a tetracycline-inducible KREH2 RNAi-plasmid construct, and Lister 427 29-13 PCF expressing a tetracycline-inducible KH2F1 RNAi-plasmid construct were grown as in our recent study (38). RNAi constructs induced with 1 µg/mL Tet (Sigma) for three days (BSF) or three to four days (PCF) downregulated the targeted protein by ∼80% in both cell stages, as reported (32,38). Growth curves and Western blotting of editing proteins determined these time points (13,38).

### Generation of BSF cells that exclusively express v5-tagged KH2F1 and that lack kDNA

Endogenous KH2F1 alleles were knocked out from BSF strain Lister 427 Single Marker (SM) cells using floxed blasticidin (BSD)- and puromycin (PAC)-Herpes Simplex virus thymidine kinase (HSVTK) drug cassettes as previously described (48,49). Correct insertion of knockout cassettes was assessed by PCR (see **Supplementary Table ST2** for a list of DNA oligonucleotide primers and plasmids). Drug cassettes were excised following each allele knockout via transient expression of Cre recombinase from pLEW100Cre_del_tetO (Addgene plasmid 24019; a gift from George Cross, The Rockefeller University, United States) and selection with ganciclovir (Invivogen, United States) as previously described (11,48). Prior to second allele knockout, the cells were also transfected with a *Not*I-linearized pEnT6+ATPaseGammaWT+3UTR construct that allows the replacement of an endogenous WT <u>ATPase</u> gamma subunit allele with a mutant allele containing the L262P mutation and selected with BSD (50). Expression of this mutant gamma ATPase (MGA) allele was previously shown to compensate for the loss of the mitochondrial genome and gene expression in BF *T. brucei*, including of RNA editing and editing proteins (11,50–52). Finally, the cells were transfected with *NotI*-linearized pHD1344tub(PAC)-KH2F1-Cterm3V5 plasmid, which allows for constitutive expression of C-terminal 3xV5 tagged KH2F1 from the β-tubulin locus (53). kDNA was removed from the KH2F1 null, KH2F1-v5 BSF cells that contained the MGA allele by treatment with 20 nM acriflavine, maintaining cell density between 1 × 10^5^ cells/mL and 2 × 10^6^ cells/mL (50).

### *In vitro*-induced differentiation of monomorphic BSF cells into PCF cells

*In vitro*-induced differentiation of monomorphic Lister 427 SM BSF or pleomorphic TREU 927 into replicating PCF was performed using reported conditions with some modifications (54). Cells at a density of 5 x 10^5^ cells/mL were first treated with 10 µM 8-pCPT-2’-O-Me-cAMP (aka cAMP; Cayman Chemical) in HMI-9 medium at 37°C, 5% CO2. These cells were transferred into SDM-79 medium with 3 mM each citrate/cis-aconitate to induce differentiation to replicating PCF at 26°C. Basic hallmark changes in development were verified independently in two labs using common primers for BSF and PCF-specific transcripts and pre-edited and edited mt transcripts, e.g., major surface antigens and fully edited COX2 (**Supplementary Table ST2**). Initial assays used conventional qPCR. Subsequent assays using Biomark HD Fluidigm examined a broader range of transcripts. Briefly, total RNA was harvested from ∼2 × 10^8^ cells using TRIzol pre- and two days post-addition (or not) of 10 μM cAMP, and two, four, or six days after transfer into SDM-79 medium supplemented with 3 mM each citrate/cis-aconitate. Samples labeled “d0, d2 or d8 + cAMP” in the main text figures indicate cells prior (control) or two- or eight-days post cAMP treatment, respectively. Isolated RNA was treated with 10 U of Turbo DNase (Life Technologies) according to the manufacturer’s instructions and purified through acid phenol:chloroform extraction. A 1 μg aliquot of total RNA was used to synthesize cDNA using the iScript select cDNA synthesis kit with random hexamers. cDNAs were then analyzed directly in qPCR reactions. For conventional qPCR, the cDNA was amplified using the iTAQ SYBR green supermix (Bio-Rad) and primers described in **Supplementary Table ST2** on a 96-well plate using the following thermocycling conditions: 1 cycle at 95°C for 30 s and 36 cycles of 95°C for 5 s, 55°C for 15 s and 60°C for 30 s. Reactions were conducted on a C1000 touch thermocycler plus CFX96 Real-Time System (Bio-Rad). Data were analyzed in the Bio-Rad CFX Manager real-time PCR analysis software (v3.1), using the linear (derivative) baseline correction method and the auto (global) threshold cycle (CT) method. All amplicons were cloned and verified by Sanger Sequencing, and single products were confirmed on a 2% agarose gel. For BioMark Fluidigm assays, the random hexamer generated cDNA was pre-amplified in multiplex specific-target-amplification (STA) reactions using TaqMan PreAmp master mix (Life Technologies) and with the following thermocycling conditions: 1 cycle at 95°C for 10 min and 14 cycles of 95°C for 15 s and 60°C for 4 min. Pre-amplified cDNA was treated with exonuclease I (New England Biolabs) and diluted 5- (for BSF) or 10-fold (for PCF). High-throughput real-time PCR was then conducted on the BioMark HD system with Fluidigm 48-by-48 dynamic array integrated fluidic circuits (IFCs), using SsoFast EvaGreen supermix with Low ROX (Bio-Rad) and primers described in **Supplementary Table ST2**. Primers were as reported (55) or designed in this study. Processing of the IFCs and operation of the instruments were performed according to the manufacturer’s procedures. PCR was performed using the thermal protocol GE Fast 96 × 96 PCR + Melt (v2.pcl). Data were analyzed in the Fluidigm real-time PCR analysis software, using the linear (derivative) baseline correction method and the auto (global) threshold cycle (CT) method. For both conventional qPCR and BioMark Fluidigm assays, the CT values determined were exported to Excel software for further processing. Calculations of fold changes in RNA levels in samples following *in vitro* differentiation were done via the 2 [−ΔΔC(T)] method (56) using mtRNP or TERT as an internal reference. We found similar levels of mtRNP mRNA between stages in this study. TERT mRNA was reported to be relatively constant between stages (*57*). Duplicate (BioMark) or triplicate biological replicates of each cDNA sample were generated and assayed for each target and internal reference per experiment, and C(T) data was averaged before performing the 2 [−ΔΔC(T)] calculation (56).

### Western blot and immunofluorescence analyses of *T. brucei* editing proteins

SDS-PAGE followed by Western blot analyses of BSF cell lysates to detect KREH2 and KH2F2 (REH2C complex subunits), and RESC13 (RESC complex subunit) were performed as previously described for PCF cell lysates (13,32,38). RESC2 was detected using an anti-RESC2 mouse monoclonal antibody (1:25 dilution) (58), and KH2F1-v5 was detected with rabbit anti-v5 antibody (1:5,000 dilution; Thermo Fisher), followed in each case by anti-mouse anti-IgG secondary antibody (1:5,000 dilution; Bio-Rad). Immunofluorescent microscopy of *T. brucei* BSF cells was carried out as in our recent study with some modifications (38). Dilutions for antibodies were 1:5,000 KREH2, 1:5,000 RESC1/2, and 1:1 KREL1 (i.e., 125 μL blocking buffer and 125 μL of antibody).

### Preparation of RNA for library construction, cDNA synthesis, Illumina sample preparation, and sequencing

Total mtRNA or RNA in complexes immunoprecipitated by anti-RESC6 or anti-RESC1 antibodies were isolated from four biological replicates of cells following BSF and PCF ± RNAi induction as previously described (32,38). Gene-specific cDNA synthesis was carried out with 2 µg of mtRNA using the iScript Select cDNA Synthesis Kit (BioRad) and oligo 2278 (see **Supplementary Table ST2**), which hybridizes to the 3’ terminus of ND7. We checked for the specificity of targeted cDNA synthesis by amplifying using primers containing universal Illumina adapters (1682/2608) (**Supplementary Table ST2**), cloning, and Sanger sequencing as before (32,38). Illumina libraries were prepared as previously described with modifications (13). ND7 libraries were amplified from 10 ng BSF or PCF gene-specific cDNA for 24 cycles with primers containing Illumina adapters (ND7 3’ domain 1682/2608).

### Processing RNA-seq data for ND7 3’ editing and identifying non-cognate gRNA isoforms that complement the extended 3’ element

Amplicon RNA-seq of ND7 3’ domain editing was processed as reported (13). Subsequently, sample alignment output data were further processed in the R environment (http://www.r-project.org) for summarizing and figure-generation purposes. Searches for gRNAs encoded in the *T. brucei* BSF strain EATRO 1125 minicircle genome (8) that match alternatively edited mRNA sequences were performed using Python scripts (package 3.7) as previously described (9). Alignments of predicted (annotated in minicircles) and sequenced gRNA in EATRO 1125 total mtRNA (8,9) are available online at http://hank.bio.ed.ac.uk. Alignments of sequenced gRNA in PCF strains EATRO 164 total mtRNA (10) and Lister 427 in total mtRNA and RESC6-immunoprecipitations (31) are available online at http://bioserv.mps.ohio-state.edu/RNAseq/T-brucei/MRBs/. We confirmed the sequence of pre-edited ND7 used in this study by PCR amplification of genomic DNA and Sanger sequencing of three independent amplicons. Analyses of last-edit sites were performed as follows. ND7 3’ RNA-seq data was analysed in the R-environment (v4.2.2; R core team 2022; https://www.r-project.org/) to determine the position of the last edited site in the top 100 most abundant amplicons in the corresponding RNA-seq library. Data wrangling was assisted by the “tidyverse” R package (v2.0.0; https://doi.org/10.21105/joss.01686) (59). RNA-seq amplicons were cross-referenced against the ND7 pre-edited sequence one nucleotide at a time, progressing from the 5’ to the 3’ end. The first (5’ most) detected mismatch was classified as the last edit site in that amplicon. Last-edit site frequency was calculated from the top 100 most abundant amplicons by totalling the number of reads with a specific last-edit site and dividing by the total number of reads from the 100 most abundant amplicons.

### Sample generation for *in-vitro* DMS-MaPseq and data analyses

The structure of full-length ND7 mRNA was experimentally determined *in vitro* by DMS-MaPseq (60). Synthetic gBlocks (IDT DNA Technologies) were generated containing the entire ND7 pre-edited (PE, gBlock 2404) or the most common isoform of ND7 bearing the 3’ High Frequency Element described in this study (HFE, gBlock 2405; **Supplementary Table ST2)**. gBlocks for ND7 were amplified with primers 2631 and 2632, and amplicons containing a T7 promoter were gel purified (Machery-Nagel) and cloned into plasmid pHD1344Tub(PAC) (48), creating p557 and p558, respectively. Plasmids were verified by Sanger sequencing, and linearized templates with XhoI were used for T7 *in vitro* transcription as reported for another edited mRNA (38). Briefly, 1 µg of purified DNA template was used in the Hi-Scribe T7 transcription kit (NEB) at 37°C overnight (∼16 hr.). Synthesized RNA was purified, refolded, and DMS treated prior to DMS-MaPseq library generation using IDT’s xGenTM Broad-Range RNA LibraryPrep Kit as reported with slight modifications (38). Reads were analyzed and DMS signal was determined using the DREEM algorithm (60). RNA secondary structure was predicted with the program RNAstructure v.6.0.1 (61) and visualized using VARNA v.3.93 (62).

### Isolation of *in vivo* chimeric molecules of gRNA gCOX3 with mRNA ND7 or mRNA COX3

To isolate gRNA/mRNA bimolecular chimeras *in vivo*, we generated ND7 or COX3 gene-specific cDNA using 2 µg of DNase-treated mitochondrial RNA from wild-type PCF cells. cDNA was generated using the BioRad iScript Select cDNA synthesis kit per the manufacturer’s directions with oligo 2278 (ND7) or 2732 (COX3) in a 20 µL reaction. 2 µL of resulting cDNA was used in a 50 µL Phusion HF polymerase PCR reaction using the forward oligo 2882 (COX3 gRNA) and 2278 (ND7 mRNA) or 2732 (COX3 mRNA) to enrich for chimeras. The resulting PCR product was verified on 2% agarose gel, and the corresponding bands were gel-eluted using the Nucleospin PCR cleanup kit. 10 ng of purified PCR product was used as template in subsequent PCR amplification using nested primers, which added 5’ HindIII and 3’ XhoI sites for cloning into pHD1344Tub (PAC) plasmid (2880/2881 COX3-ND7 and 2880/2883 COX3-COX3). The undigested PCR product was cloned into HindIII/XhoI digested pHD1344Tub(PAC) plasmid using the In-Fusion cloning kit at a 2:1 molar ratio. This reaction generated plasmids p658 (COX3-ND7) and p659 (COX3-COX3). 2 µL of In-Fusion product was then transformed into Stellar competent cells per the manufacturer’s instructions, plated on Ampicillin-selective LB agar plates, and allowed to grow overnight at 37°C. Ten individual colonies were picked from each plate (COX3-ND7 or COX3-COX3). Plasmid was isolated, inserts were amplified and analyzed by Sanger sequencing, and sequences were examined using the MUSCLE alignment tool as previously described (32,38).

### Calculations and statistical analysis

Total editing and NC/C (non-canonical/canonical) ratio values (previously termed Inc/Cor ratio) of the percentage of reads, both site-by-site and cumulative, were calculated as previously reported (13,38). This updated nomenclature reflects a key observation here and in a recent study that specialized non-canonical editing (38) and canonical editing are developmentally controlled by KREH2. Each site exhibits an NC/C edit ratio <1 since the NC edits percentage (numerator) is smaller than C edits. In plots of cumulative NC/C along the fragment, added-up individual NC/C increments (each <1) generate a cumulative value >1. Graphs compare replicate sets for two conditions of either total mtRNA or RESC6- or RESC1-RNA immunoprecipitations (RIPs) (e.g., PCF vs. BSF or -Tet vs. +Tet), where one replicate set includes at least three biological replicates, and another set includes at least two biological replicates. These replicate sets enabled statistical calculation of p-values, average, and standard deviation. A description of samples and p-values for all sets compared are included (**Supplementary Table ST3).** Significant changes in editing fidelity (NC/C value) in each site were determined as before (13,38). To generate *P* values for the effects of KH2F1-RNAi on ND7 3’ editing, we combined independent biological replicates for 3- and 4-days post-induction of RNAi since these two type points of treatment had comparable outcomes. This allowed us to increase the total number of replicates for the +Tet condition. We used one-way ANOVA to test the null hypothesis that there is no significant difference between groups, with this null hypothesis rejected at *P* < 0.05. The mean + SD of independent biological replicates was reported.

## RESULTS

### KREH2 localizes near kDNA and associates with KH2F1 and KH2F2 in REH2C complexes of BSF cells

Most editing studies have been performed in PCF cells because they grow at higher densities and generate more material than BSF cells. Our prior studies showed that KREH2 is essential for growth and editing in both PCF and BSF *T. brucei* (38,41,43). We also showed that KH2F1 (REH2C) co-localizes with kDNA (38) and confirmed KREL1 (RECC) and RESC1/2 (RESC) co-localization with kDNA (43,63). We now show for the first time that KREH2 co-localizes near or with kDNA in BSF cells (**Fig. 1A; Supplementary Fig. S1A**). Furthermore, KREH2 interacts in an RNA-independent fashion with KH2F1 and KH2F2 in a BSF dyskinetoplastic (DK) mutant strain that lacks kDNA and is thereby devoid of mtRNA (50) (**Fig. 1B; Supplementary Fig. S1B**). Surprisingly, the steady-state levels of KH2F2, RESC2, and RESC13 evidently decreased in mt-extract from DK cells. Finding this effect on proteins in RESC and REH2C in the absence of mtRNA is unprecedented and implies an induced-fit mechanism in RNA-protein recognition that impacts protein stability during assembly of these RNPs (64,65). This observation is in line with RESC and REH2C forming stable RNPs. In contrast, the stability of RECCs, which transiently interact with RNA, is not affected in DK cells (66). To our knowledge, this is the first time that RESC or KREH2C components have been compared in WT vs DK versions of the same strain. This novel finding is not further addressed here and will require a separate study. Thus, RNA-free versions of the REH2C complex exist in BSF as originally proposed in PCF (32).

**Figure 1.**
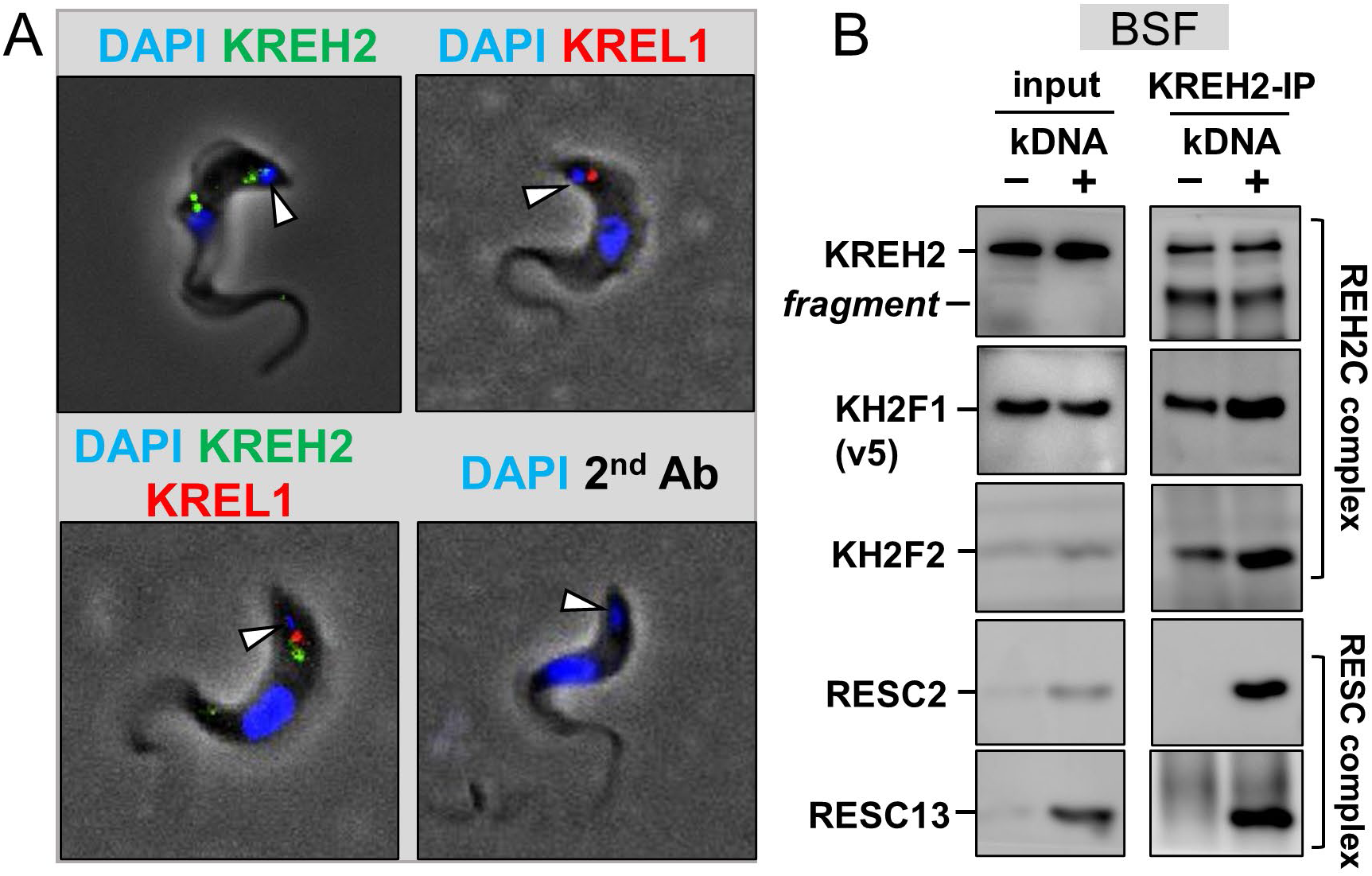
KREH2 localization and association with proteins in REH2C in BSF cells. **(A)** Immunofluorescent microscopy of BSF *T. brucei*. Cells were imaged for marker proteins in editing complexes: KREH2 (REH2C) and KREL1 (RECC). White arrows point to DAPI-stained kDNA in each cell. **(B)** Western blot of mt-extracts (input) and KREH2 immunoprecipitation (KREH2-IP) from a BSF dyskinetoplastic cell line that we made via acriflavine treatment as described in the methods section, which is devoid of kDNA (-kDNA). The parental cell line that was not treated with acriflavine and contains kDNA (+kDNA) was included as a control. These BSF cell lines are conditional nulls of KH2F1 that constitutively express KH2F1-v5 tagged protein, as described in the methods section. KREH2, KH2F1-v5, KH2F2 (all in REH2C), and RESC2 and RESC13 (in RESC) were examined. KREH2 is often fragmented (*fragment*) in our mitochondrial extract preparations.

### KREH2 knockdown differentially affects total ND7 3’ domain editing in PCF and BSF

Prior qRT-PCR assays confirmed that KREH2 promotes accumulation of mature fully edited mRNAs in PCF and BSF *T. brucei* (38,41,43). However, these assays do not inform the precise details of 3’-5’ editing progression. Here, we applied targeted base-resolution RNA-Seq to examine how KREH2 knockdown in PCF and BSF may affect the editing progression of the 3’ terminus in ND7 3’ domain (aka ND7 3’), including ORF (90 sites) and UTR (7 sites) sequences in the amplified fragment (**Supplementary Fig. S2A**). We began by examining ND7 3’ in total mtRNA. To illustrate the raw data initially collected and tallied by our bioinformatics pipeline, we included a stack plot of representative biological replicates in PCF and BSF cells (**Figs. 2A-B**). These plots provide a snapshot of all editing events scored at each site in the ND7 3’ amplicon.

**Figure 2.**
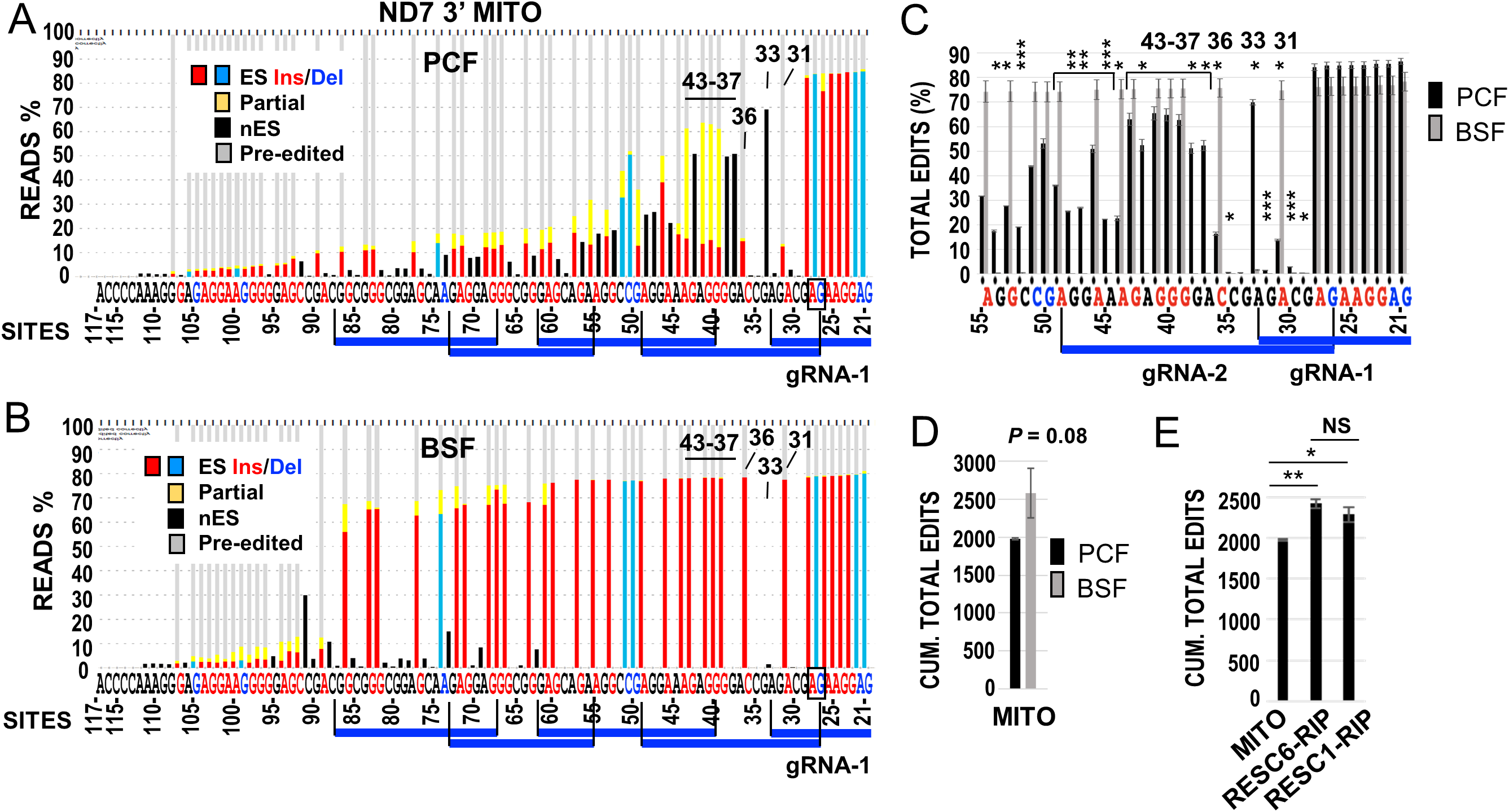
Comparison of ND7 3’ total editing in PCF and BSF cells. ‘Snapshots’ of typical ND7 3’ amplicon RNA-seq data sets from **(A)** PCF and **(B)** BSF. Stacked histograms show all possible editing events at each site in replicate mtRNA (Mito) samples from cells containing a KREH2-RNAi construct -Tet (i.e., uninduced), as in all other panels below, which used the same uninduced construct. Color-coded nucleotides are just 3’ to canonical sites for U-insertion (Ins, red), U-deletion (Del, blue), or sites not expected to change in mature mRNA (black) (see **Supplementary Table S1**; glossary of terms). Bars represent the percentage of canonical insertion (red), deletion (blue), or non-canonical edits (yellow) at canonical sites or edits at sites not expected to change (black). Blocks of canonical editing thought to be directed by known gRNAs are indicated by indigo lines. **(C)** Site-by-site analysis of total edits across gRNA-1 and gRNA-2 (through site 55) in mtRNA from PCF vs. BSF cells. **(D)** Cumulative total edits in PCF vs BSF mtRNA. **(E)** Cumulative total edits from mtRNA, RESC6-RNA-Immunoprecipitation (RIP), and RESC1-RIP. The cumulative value at the most 5’ site (site 117) in the amplicon was plotted. Full-amplicon analyses are in **Supplementary Fig. S2C**. As noted above, all cells in this figure are uninduced. Average and error bars representing the standard deviation of biological replicates to determine *P*-values *** < 0.005, ** < 0.05, * < 0.5 were annotated. See Material and Methods and **Supplementary Table S3** for additional details on statistical analysis.

The total editing action on an mRNA comprises two separate components, canonical and non-canonical. Changes between these two types of edits were not necessarily reciprocal in previously examined A6 and RPS12 mRNA substrates (13,38). We focused on the 3’ region of ND7 3’ because our plots exhibited extensive editing action and dramatic differences along the first few blocks between BSF and PCF (**Figs. 2A-B**). The first gRNA, initiator gRNA-1, covers most of the 3’ UTR and also two sites that create the stop codon. Editing is highly efficient along these first few sites in both PCF and BSF. However, the region spanned by the 3’ terminus of gRNA-1 (e.g., sites 31 and 33) and subsequent guides exhibited evident differential changes. In BSF, editing action, mostly canonical, remained high along the first five gRNAs but dropped dramatically near the 5’ end of the examined fragment. In contrast, in PCF, canonical editing suddenly dropped at site 31 and stayed low in the remaining examined fragment, whereas partial non-canonical editing (i.e., yellow and black bars) was substantial, particularly opposing the 3’ end of gRNA-1 (at site 33) and across gRNA-2. We hypothesized that early editing across the first two guides might offer early differential checkpoints in ND7 maturation during development, which we examined in more detail.

In the ND7 3’ fragment amplicon, the forward primer matched a pre-edited sequence, which likely selects for the low editing action at the 5’ end of the examined sequences. Because the mRNA 3’ terminus is short and AU-rich, a suitable reverse primer was used to tally U-indels beginning at the second position for canonical editing (site 21) in ND7 3’ (**Supplementary Fig. S2A**). The initiator gRNA-1 in our alignments represents gRNA-1 gND7(1269–1320) (22), aka gND7 *B1* (*31*) in PCF strains EATRO164 and Lister 427, respectively. gRNA-2 in our alignments is one of two reported potential guides, gND7(1240–1268) (22), aka gND7 *B2.alt* (*31*) in PCF strains EATRO164 and Lister 427, respectively. We reported that gND7 *B2.alt* best matches fully-edited ND7 3’ examined by Sanger-sequencing in PCF strain Lister 427 (31,42). These first two guides also generate the best match with the ND7 3’ canonical pattern in our BSF samples. In BSF strain EATRO1125, the annotated guides in minicircle DNA libraries predict the same ORF in ND7 3’ except that it ends in VDR (9) (**Supplementary Fig. S2B**). This is one of two alternative ORFs proposed in EATRO164 ending in VDR or EYR (22). The alternative sequence forming EYR was present in up to ∼70% and ∼14% of all reads in BSF and PCF in Lister 427 in this study, in line with our previous sequencing studies (31,42). The alternative sequence forming VDR was present in 0.2-1% of all reads in our samples, and its frequency was not evidently affected by cell stage or KREH2 expression (see the statistics in **Supplementary Table ST3**). Thus, canonically edited ND7 3’ with a predicted ORF ending in EYR may be the “functional version” in strain Lister 427 under control during development.

All our subsequent analyses directly compared independent biological replicates of each sample plus or minus KREH2 knockdown. We focused on a region that spans the first two PCF gRNAs in the ND7 3’ editing domain, particularly sites 33 and 37-to-43, which includes extensive non-canonical editing. However, analyses of entire amplicons were also included (**Supplementary Figs. S3-S7**). We plotted total editing at each site as well as cumulative editing, either along the amplicon or up to the most 5’ site examined. We compared total mtRNA in PCF and BSF and RNA-immunoprecipitations (RIPs) of RESC6 and RESC1 in PCF. Among many potential RESC isoforms (30,31,67), RESC-A contains RESC6 and RESC1, while RESC-B contains RESC6 but not RESC1 (36). This is the first time isolated native RESC1 complexes have been examined by RNA-Seq, as was previously done for native RESC6 complexes (13,38). Total mtRNA samples without KREH2 knockdown confirmed significant differences between PCF and BSF in analyses of total editing action at most sites examined (**Fig. 2C; Supplementary Fig. S2C**). As expected, cumulative total editing at the most 5’ site exhibited higher editing action in BSF than PCF (**Fig. 2D**). Also, total editing action was significantly enriched in both RESC6 and RESC1 RIPs vs. total mtRNA (**Fig. 2E**). This was expected since our earlier RT-qPCR studies had shown enrichment of fully-edited and pre-edited ND7 3’ in RESC6-RIPs vs. total mtRNA (31).

KREH2 knockdown in PCF (32,38) significantly decreased total editing in mtRNA and RESC6-RIP samples. This effect was observed in most sites examined (**Figs. 3A-B; Supplementary Figs. S3A-C**) and in graphs of cumulative total editing up to the most 5’ site (**Figs. 3C-D**). PCF knockdown of KH2F1, another subunit of REH2C, confirmed the above effect on total editing in mtRNA and RESC6-RIPs (**Figs. 3C-D**; **Supplementary Fig. S3D**). We previously reported that KH2F1 knockdown in PCF destabilizes KREH2, effectively creating a dual knockdown (32). Thus, specific KREH2 knockdown, which does not affect the KH2F1 integrity (32), suffices to decrease ND7 3’ total editing in PCF. Cumulative total edits were similar in RESC1 and RESC6-RIPs, but total editing inhibition by KREH2 knockdown was more robust in RESC1-bound complexes (**Fig. 3D)**. KREH2 knockdown in BSF also generally decreased total editing in mtRNA in the sites examined (**Supplementary Figs. S3B**). These results are in line with prior qRT-PCR assays showing that REH2C loss-of-function reduced the steady-state accumulation of mature fully edited ND7 3’ in PCF and BSF (13,32,38,42).

**Figure 3.**
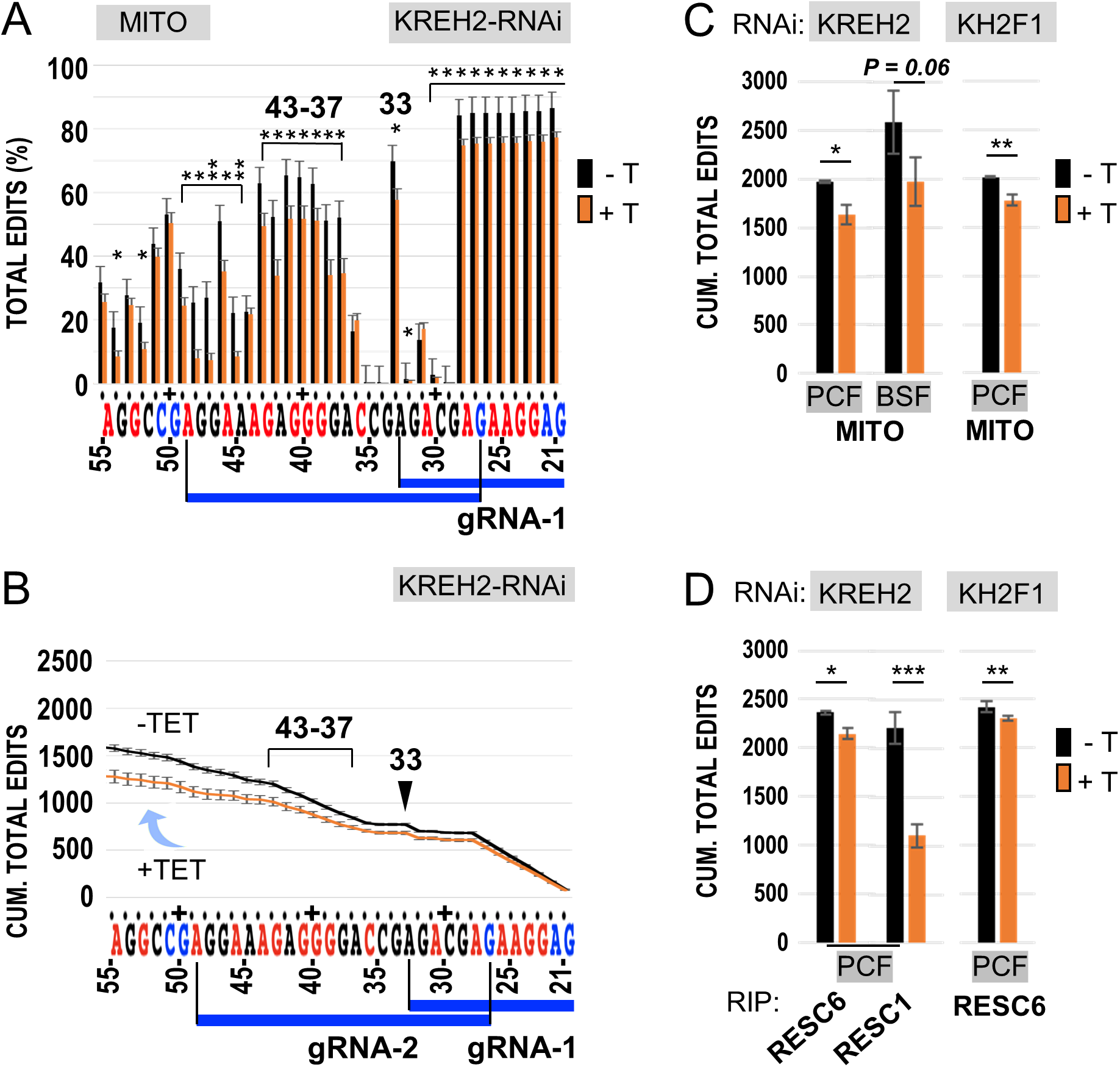
Analyses of ND7 3’ editing domain total editing upon KREH2-RNAi and KH2F1-RNAi in PCF and KREH2-RNAi in BSF. **(A)** Site-by-site, and **(B)** cumulative total edits across gRNA-1 and gRNA-2 (through site 55) in PCF mtRNA ± KREH2-RNAi. RNAi knockdowns decreased the targeted protein by ∼80% (see methods section). Sites 33 and 37-43 of particular interest in ND7 3’ early editing, as described in the text, are highlighted. **(C)** Cumulative total edits up to the most 5’ site (site 117) in the ND7 3’ domain amplicon in PCF mtRNA ± KREH2-RNAi or KH2F1-RNAi and BSF mtRNA ± KREH2-RNAi. **(D)** Cumulative total edits as in panel C but in RESC6- and RESC1-RIPs in PCF KREH2-RNAi or KH2F1-RNAi. Full-amplicon analyses are in **Supplementary Fig. S3**. Black bars are +Tet, and gold bars - Tet (labeled ± T). Statistical analyses are as in Fig. 2.

### KREH2 knockdown differentially affects ND7 3’ editing fidelity and pausing in PCF and BSF

We asked whether KREH2 affects editing fidelity in ND7 3’, particularly across the first two guides, where dramatic differences in editing action were apparent between PCF and BSF. Large NC/C values indicate low editing fidelity in site-by-site and cumulative plots. As mentioned above, the canonical sequence examined here likely represents a functional ND7 3’.

Remarkably, KREH2 or KH2F1 knockdowns in PCF increased editing fidelity (i.e., significantly reduced the NC/C ratio) in the ND7 3’ edited domain. This effect was observed at most sites examined, starting at sites directed gRNA-1, in site-by-site and cumulative NC/C plots in mtRNA and RESC6-RIPs (**Figs. 4A-D; Supplementary Figs. S4A-D; Figs. S5A-C)**. Generally, increased editing fidelity in the knockdowns involved concurrent upregulation of canonical edits and downregulation of non-canonical edits (**Figs. 4E-F; Supplementary Figs. S6-S7).** As expected, RESC6-RIPs exhibited higher cumulative canonical and non-canonical editing than mtRNA (*P*= 0.01 and *P*= 0.0007, respectively). Regulation was particularly dramatic in non-canonical edits including sites 33 and 37-43 in both mtRNA and RESC6-RIPs (**Supplementary Figs. S6**). These results suggest that REH2C actively represses ND7 3’ domain editing in functional editosomes.

**Figure 4.**
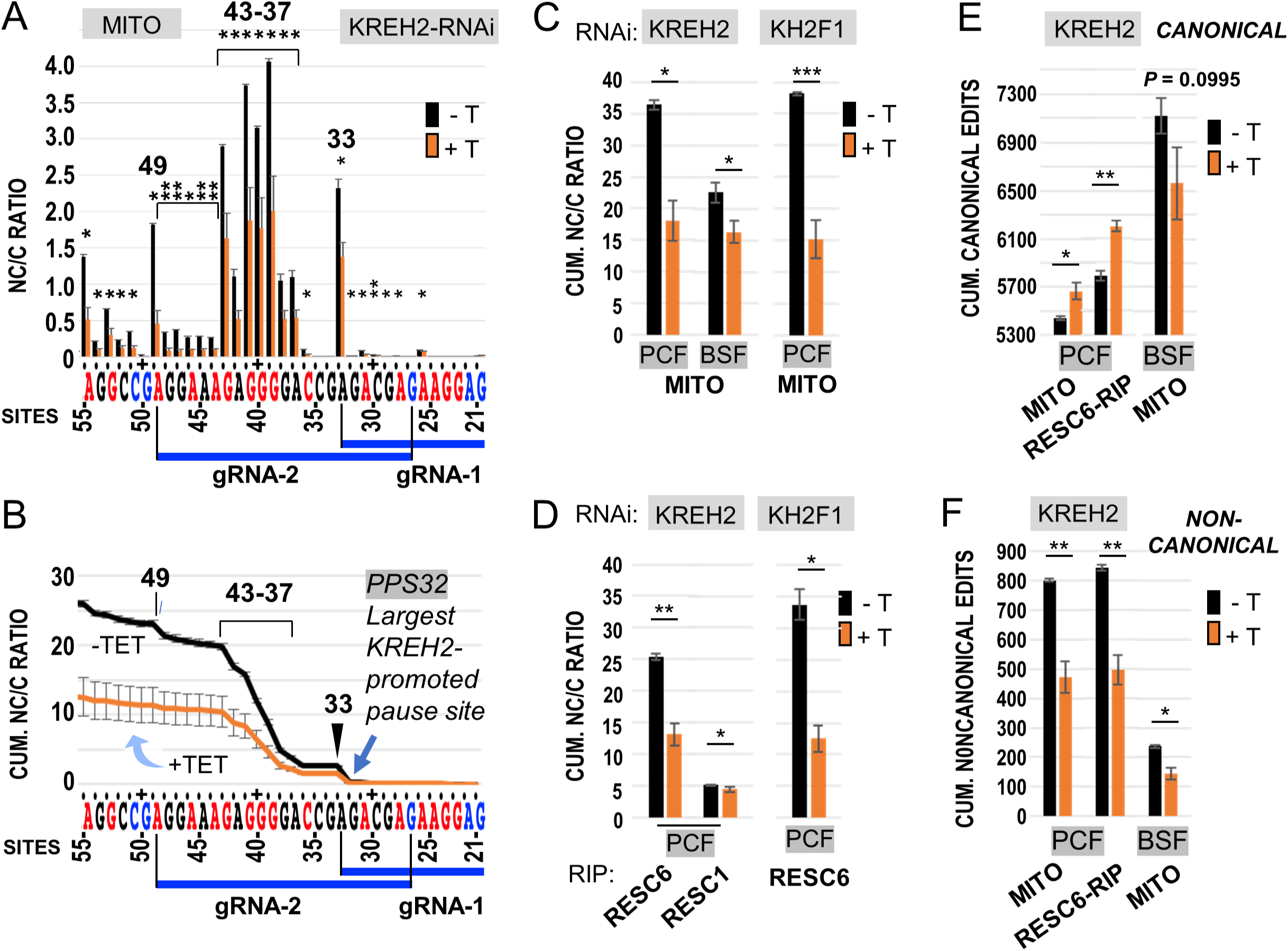
Analyses of ND7 3’ domain editing fidelity (NC/C) upon KREH2-RNAi or KH2F1-RNAi in mtRNA or RESC. **(A)** Site-by-site and **(B)** cumulative NC/C ratios across gRNA blocks 1-2 in PCF mtRNA ± KREH2-RNAi. NC/C ratios are the percentage of non-canonical reads divided by the percentage of canonical reads at the same site. Sites 33, 37-43, and 49 (highlighted) exhibited particularly high NC/C ratios in the replicates. **(C)** Cumulative NC/C ratios at the most 5’ site (site 117) in the ND7 3’ domain amplicon in PCF mtRNA ± KREH2-RNAi or KH2F1-RNAi and BSF mtRNA ± KREH2-RNAi. **(D)** Cumulative NC/C ratios as in panel C but in RESC6-and RESC1-RIPs in PCF ± KREH2-RNAi or KH2F1-RNAi. **(E)** Cumulative canonical and **(F)** cumulative non-canonical edits at the most 5’ site (site 117) in mtRNA and RESC6-RIP in PCF or BSF ± KREH2-RNAi. All RNAi is ± Tet (labeled ± T). Note that the +Tet decreased NC/C, i.e., increased editing fidelity. Full-amplicon analyses are in **Supplementary Fig. S4** and **Table ST3.** The highest NC/C in the amplicon at site 33 (arrowhead) caused the largest KREH2-promoted pause at site 32 (PPS32) in the sequence examined. Other annotations and statistical analyses are as in Figs. 2 and 3.

In other comparisons, cumulative NC/C in ND7 3’ was higher in mtRNA (∼five-fold) and RESC6-RIPs (∼twenty-fold) vs. RESC1-RIPs (**Fig. 4D)**. However, RESC1-RIPs still exhibited a small, but statistically significant increase (*P=* 0.01) in ND7 3’ editing fidelity in the KREH2 knockdown. Thus, RESC6-containing RESCs seem a major platform for KREH2-dependent ND7 3’ editing repression in PCF. Also, although ND7 3’ editing fidelity was generally higher in BSF than PCF, KREH2 knockdown decreased the NC/C ratio at several sites in BSF. However, knockdown of KREH2 in BSF did not upregulate canonical edits as in PCF. In BSF, the percentage of non-canonical editing is low, but canonical editing is high; upon KREH2 knockdown the former decreased significantly, but the latter declined moderately (P ∼0.1) (**Figs. 4E-F)**. Combined, these changes reduced the cumulative NC/C ratio in BSF (**Fig. 4C)**. Importantly, KREH2 knockdown in BSF did not significantly affect the NC/C ratio at sites 33 and 37-43 **(Supplementary Figs. S4B, S4E)**.

Sites where canonical editing naturally decreases are known as intrinsic pause sites (IPSs) (12). Peaks in NC/C indicate pausing in canonical editing from one site to the next. That is an interruption in canonical progression due to persisting alternative edits immediately upstream (13). In PCF, IPS32 was the largest pause site identified in the ND7 3’ fragment examined, i.e., NC/C at site 33 was 107-fold and 160-fold higher than at site 32, in RESC6-RIPs and mtRNA, respectively (**Figs. 4A-B; Supplementary Figs. S4)**. Thus, canonical editing may continue past IPS32 in ∼1 out of 107 or 160 molecules in RESC-RIPs and mtRNA, respectively. As mentioned above, the KREH2 knockdown reduced the NC/C at most sites examined, including site 33, thereby alleviating IPS32 to facilitate editing progression. In typical IPSs, canonical editing resumes at the following upstream site. However, we observed tracks of sites, each site exhibiting high NC/C, that is, an “intrinsic pause track,” including sites 37-43. KREH2 knockdown alleviated this and other intrinsic pause tracks in ND7 3’ in mtRNA and RESC6-RIP samples. Among others, particularly high NC/C values in ND7 3’ identified IPS35 and IPS36 next to the 37-43 site track in block 2 (by gRNA-2), and IPS48 and IPS54, next to sites 49 and 55, respectively (**Figs. 4A-B; Supplementary Fig. S4)**. Major IPS stimulated by KREH2 were termed KREH2-promoted pause sites (PPSs). As indicated above, IPS32 may be the largest KREH2-PPS in ND7 3’ in PCF.

In brief, PCF-specific repression of ND7 3’ mRNA editing involves decreased editing fidelity and increased pausing in this sequence. This repression requires KREH2, is effective in native RESC6-bound RESC complexes, and particularly affects early editing in PCF.

### Opposite modulation of non-canonical and canonical editing by KREH2 reveals a major early checkpoint in PCF

To further examine the source of ND7 3’ low editing fidelity (high NC/C ratios) in PCF, we surveyed all editing events, particularly at sites 33 and 37-43, near the ORF 3’ end (**Figs. 5A-B**). Analyses of all non-canonical reads revealed dominant events at these sites, defined here as specific high-frequency events (at least 75% of all NC reads) at a given site. For example, at site 33, a +2U insertion event in RESC6-RIPs accounted for 94% of the NC value (∼60% of all reads, i.e., canonical, non-canonical, and pre-edited). Dominant non-canonical reads at sites 37-42 accounted for ∼80 to 90% of the NC value (>40% of all reads) and slightly less at site 43 (**Fig. 5C**). These non-canonical reads were also present in RESC1-RIPs but less frequent (<10% of the NC value; < 5% relative to all reads) (**Fig. 5C**). Interestingly, the +2U insertion at site 33 might be directed by two encoded 3′-terminal adenines in gRNA-1 (**Fig. 5B**). These adenines are not part of the classified canonical guiding domain in gRNA-1 but are conserved in the sequenced gRNA-1 in strains Lister 427 (used in this study), EATRO 164, and EATRO 1125 (**Supplementary Fig. S2B**) (9,22,31). We previously reported a dominant non-canonical +2U insertion in mRNA RPS12 potentially directed by two conserved 3′- terminal adenines of RPS12 gRNA-1 (13). A +2U insertion by conserved 3′-terminal adenines in gRNA-1 in distinct substrates suggests that this event is biologically important. We also noted a second non-canonical event potentially directed by the 3’ terminus of gRNA-1. Such an event would create unedited site 31, potentially affecting gRNA-2 anchor binding (**Figs. 5A-B**). This additional non-canonical event may contribute to the sudden drop of canonical editing at site 31 in PCF (**Fig. 2A**).

**Figure 5.**
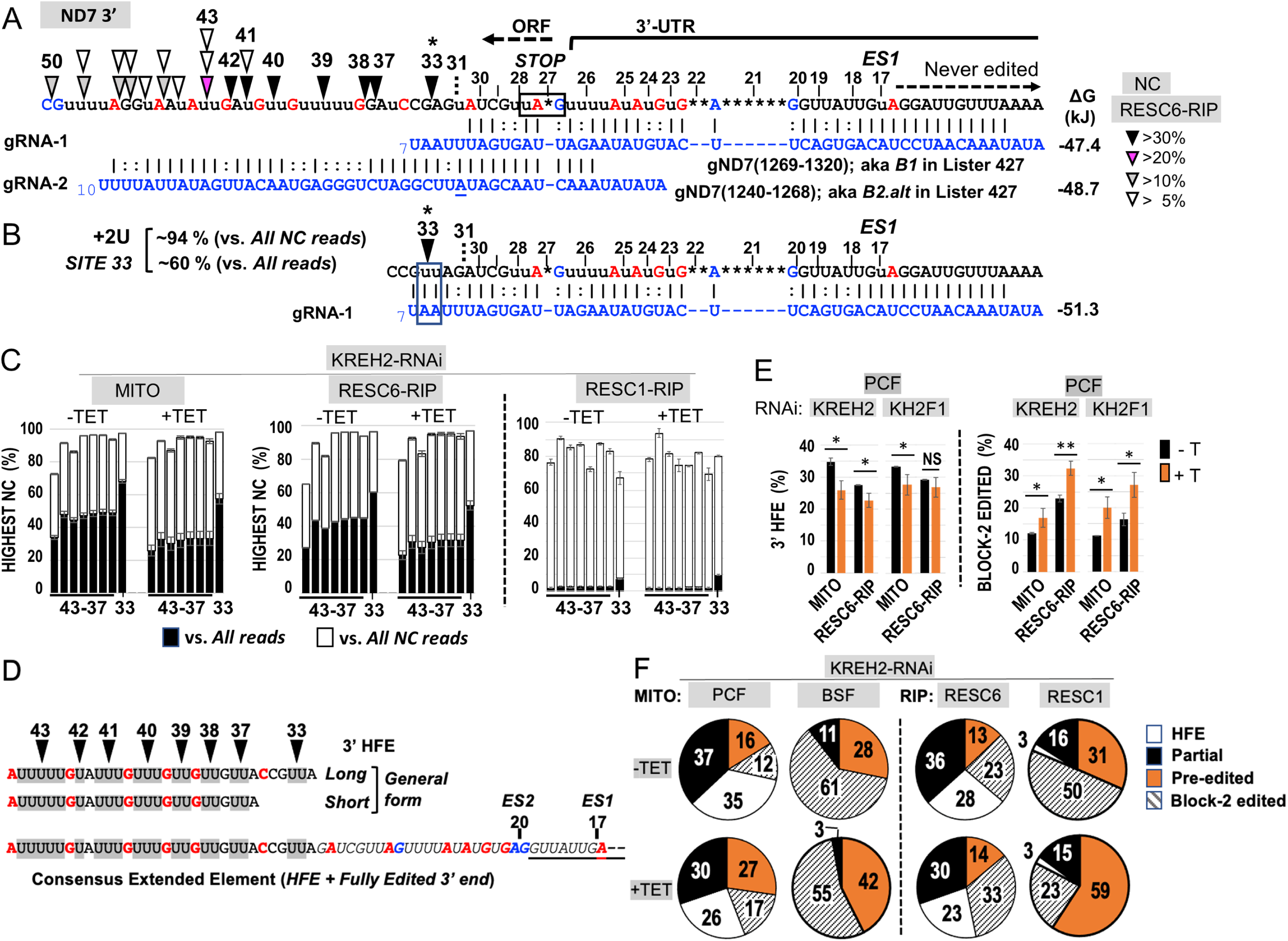
An abundant non-canonical 3’ high-frequency element (HFE) sequence in the ND7 3’ editing domain in PCF and BSF, and its modulation by REH2C proteins. **(A)** Canonically edited ND7 3’ terminus. Color-coded letters are just 3’ of sites requiring insertion (red), deletion (blue), or no changes (black). ORF, 3’ UTR, and never-edited regions are indicated. The first editing site (*ES1*) is at position 17, counting from the 3’ end. Illumina sequenced gRNA isoforms: gRNA-1 *B1* and gRNA-2 *B2.alt* in strain Lister 427 (31) correspond to gRNA-1 gND7(1269–1320) and gRNA-2 gND7(1240–1268) in strain EATRO 164 (22), with identical guiding function at block 1 and block 2, respectively. These guides produced the best match with the ND7 3’ edited pattern in this study and prior Sanger sequencing in our strain (31). Color-coded arrowheads indicate sites with a dominant NC read representing >30% (black) or less in the color-coded scale of all reads in RESC6-RIPs in uninduced (-Tet) PCF cells containing a KREH2-RNAi construct. **(B)** Canonical and non-canonical guiding potential of the initiator gRNA-1 *B1*. Alternative uninterrupted alignment of the 3’ terminus in gRNA-1 would involve non-canonical addition of +2 U at site 33 and 0 U at site 31. **(C)** Actual percentage of the dominant NC read at each indicated site versus all reads (black) or versus all NC reads (white) in mtRNA and RESC6 or RESC1-RIPs in PCF. **(D)** Consensus 3’ High-Frequency Element (3’ HFE) made by the dominant NC reads at sites 33-43. The top two 3’ HFE isoforms found in all samples examined show the dominant NC reads in gray. General 3’ HFE long or short forms, where sites 34-36 included any T nucleotide number. Bottom: ∼62-nt extended 3’ element, including the 3’ HFE and 3’ terminal fully edited sequence. (E) Frequency of 3’ HFE (*left*) or canonically edited block 2 (*right*) in mtRNA or RESC6-RIPs in PCF ± KREH2-RNAi or KH2F1-RNAi. **(F)** Frequency of 3’ HFE, and either block 2 read type: canonically edited, pre-edited, and other NC (“partial”) in PCF and BSF ± KREH2-RNAi cells. ±Tet (also labeled ±T). Statistical analyses are as in Figs. 2 and 3.

The proportion of dominant non-canonical edits at the cluster of sites that include 33 and 37-43 was particularly high but decreased in RESC6-RIPs (*P=* 0.006) and mtRNA (*P=* 0.02) upon KREH2 knockdown. While the +2U event at site 33 exhibited by far the highest percentage of non-canonical reads in ND7 3’, we hypothesized that dominant non-canonical reads at sites 33 and 37-43 may co-exist in a sequence element in ND7 3’ (**Supplementary Fig. S6)**. To test this idea, we searched for a consensus element in two versions, including sites 33-43 (long) and 37-43 (short) (**Fig. 5D**). Since sites 34-36 have variable U content, they were allowed any U number. These searches revealed a 3’ High-Frequency Element (3’ HFE) in ∼30 % of ND7 3’ amplicons in all PCF samples without KREH2 knockdown (**Fig. 5E**, left panels -Tet). The long and short elements exhibited very similar frequencies. Over 99% of the hits with the short form contained the long form. This suggested that the dominant non-canonical events at sites 33 and 37-43 were concerted.

We then asked whether KREH2 affects the frequency of the 3’ HFE in the ND7 3’ editing domain. Knockdown of KREH2 in PCF significantly decreased the 3’ HFE reads in mtRNA and RESC6-RIPs (**Fig. 5E**, left panels). KH2F1 knockdown in PCF confirmed a decrease in 3’ HFE reads in mtRNA. In RESC6-RIPs, the average 3’ HFE reads appeared to decrease upon KH2F1 knockdown, but this change was not significant, presumably due to variation between our KH2F1-RNAi sample replicates. Because the 3’ HFE is installed in the same region for canonical editing by gRNA-2, we asked whether 3’ HFE formation affects block 2 maturation. Remarkably, knockdown of KREH2 or KH2F1 significantly increased block 2 maturation in PCF mtRNA and RESC6-RIPs (**Fig. 5E**, right panels). To our knowledge, KREH2 is the first identified protein that specifically represses the maturation of an entire editing block in any substrate. Other blocks were examined below. KH2F1 may participate at least by affecting KREH2 stability in PCF (32).

To better understand how KREH2 hinders block 2 maturation in PCF, we compared the percentage of 3’ HFE, canonical, pre-edited, and remaining non-canonical “partial” editing reads in the block 2 region (**Fig. 5F**). The pool of partial reads may include related 3’ HFE variants that did not exactly match the consensus element in our searches, plus unrelated sequences of unclear origin. We calculated the percentage of partial reads by subtracting the total reads minus the sum of other read types in the block 2 region: pre-edited, canonical, and consensus 3’ HFE. In RESC6-RIPs in PCF, pre-edited reads were the lowest fraction (13%), consistent with these samples representing active RESC variants. Among read types exhibiting any editing action, partial edits were the highest (36%), followed by 3’ HFE (28%) and canonical reads (23%). Upon KREH2 knockdown, canonical reads became the highest (33%) and 3’ HFE the lowest (23%). Partial reads decreased (30%), while pre-edited reads were minimally affected, remaining the lowest (14%) in the knockdown. Thus, during early ND7 3’ editing in native RESC6 complexes, KREH2 mostly affected the ratio of edited product types (non-canonical vs. canonical sequence) rather than the overall amount of substrate consumption. In total mtRNA in PCF, KREH2 knockdown also decreased 3’ HFE and partial reads while increasing canonical and pre-edited reads. In RESC1-RIPs in PCF, 3’ HFE reads were rare (3%) and minimally affected upon KREH2 knockdown. Notably, in BSF, the presence of the 3’ HFE was negligible in mtRNA (0.01%), consistent with the previously observed efficient ND7 3’ maturation in this lifecycle stage. Thus, the 3’ HFE was, on average, ∼3,000-fold enriched in PCF vs. BSF in ND7 3’ amplicons from mtRNA. We also asked whether KREH2 knockdown promotes the maturation of other editing blocks in ND7 3’. Importantly, in RESC6-IPs, KREH2 knockdown significantly increased the maturation of blocks 3 and 4 but not block 1 (**Fig 6A)**. We found similar results in mtRNA samples, except that block 1 maturation decreased upon KREH2 knockdown. Our results in the examined blocks 2 to 4 align with the global observed changes in the NC/C ratio and the individual NC and C values at most sites in ND7 3’ (**Supplementary Figs. S6-S7)**. Thus, KREH2 may modulate the selection of gRNAs, cognate vs. non-cognate, along most of the ND7 3’ editing domain sequence examined specifically in PCF.

**Figure 6.**
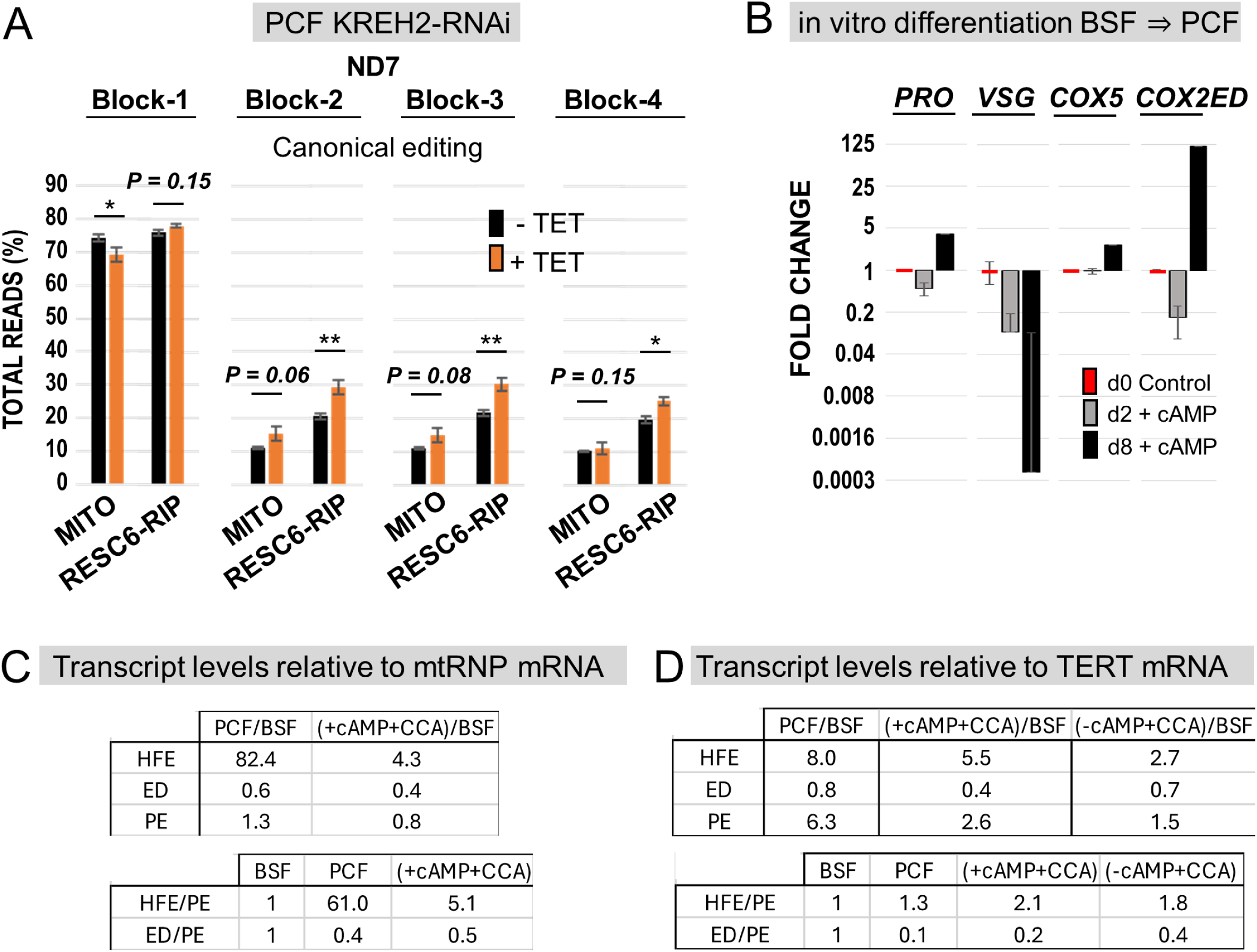
Relative changes in ND7 3’ edited blocks 1-4 upon KREH2-RNAi knockdown and in 3’ HFE upon *in vitro* differentiation. **(A)** RNA-seq analyses of the total reads (%) of canonically edited ND7 3’ domain blocks 1 through 4 in mtRNA or RESC6-RIPs in PCF ± KREH2-RNAi ±Tet. **(B)** qPCR analyses of relative levels for hallmark transcripts during *in vitro* differentiation. BSF strain Lister 427 cells before (day 0; d0) and after (day 2, d2, or day 8, d8) *in vitro* differentiation to PCF via 2 days of incubation at 37°C in BSF media +pCPTcAMP (cAMP) and 6 more days of incubation in PCF media with citrate/cis-aconitate at 26°C (full details of in vitro differentiation protocol in Materials and Methods). **(C-D)** Independent *in vitro* differentiation experiments and qPCR analyses (from two different labs; see Materials and Methods for more details) of 3’ HFE, canonically edited, and pre-edited ND7 normalized to mt-RNA polymerase mRNA (C) or TERT (D) as reference transcripts. Ratios in parent cells, PCF/BSF, or post/prior differentiation (±cAMP +CCA)/BSF are compared for the independent experiments in Lister 427 cells grown separately in two different labs for decades.

These studies so far have compared PCF vs. BSF (monomorphic) cell lines of strain Lister 427. However, these lines have been cultured separately for over a decade. A central question is whether the 3’ HFE accumulates in PCF during development. We addressed this question using *in vitro-*induced differentiation of cultured BSF strain Lister 427 monomorphic and strain TREU 927 pleomorphic cells into replicating PCF using reported conditions (54). Conventional RT-qPCR showed the expected changes of major surface antigens during differentiation, i.e., increased procyclin, and massive loss of variant surface glycoprotein mRNAs (54). We also confirmed a robust PCF-specific accumulation of fully-edited COX2 mt mRNA (i.e., activation of oxidative phosphorylation) (**Fig. 6B)** (68). These and additional changes in a broader set of transcripts during differentiation were confirmed by Biomark Fluidigm assays, including a robust increase of fully-edited COX3, and CYb transcripts, and a moderate increase of nuclearly encoded COX5, COX6, and COX10 (**Supplementary Fig. S8A**). To assess 3’ HFE accumulation in these cells, we established an RT-qPCR assay to quantitate this element relative to canonically edited and pre-edited ND7 3’ domain (**Figs. 6C-D**; **Supplementary Fig. S8B-C**). We conducted independent differentiation experiments in two labs and scored transcripts by qPCR normalized to mt-RNA polymerase (mt-RNP) mRNA or telomerase reverse transcriptase transcript (TERT), which we found (mt-RNAP) or was reported (TERT) to be relatively stable between *T. brucei* lifecycle stages (57).

Notably, these independent RT-qPCR measurements confirmed relatively higher or lower 3’ HFE and ND7 3’ canonical editing in parental PCF vs. BSF cells, respectively, regardless of the strain tested. *In vitro* differentiated PCF (±cAMP + CCA conditions) exhibited partially increased 3’ HFE and decreased ND7 3’ canonical editing in both experiments. Further normalization of these transcripts to pre-edited substrate, which was elevated in PCF vs. BSF, also confirmed the expected changes in 3’ HFE and ND7 3’ canonical editing upon differentiation.

Thus, *in vitro* differentiation recapitulated the opposite changes in 3’ HFE and canonical editing observed in PCF vs. BSF parental cells and indicated that these changes represent a bonafide phenomenon during development. We note that the detected changes in these transcripts by RT-qPCR were generally more modest than those captured in Ilumina samples. This discrepancy between the two quantitative approaches may partly reflect differences in primer binding efficiency for the amplicons examined in our RT-qPCR conditions.

Overall, PCF-specific repression of ND7 3’ involves KREH2-mediated opposite modulation of canonical and non-canonical editing in entire blocks. At a major checkpoint in early editing, repression involves inhibition of canonical block 2 editing and abundant 3’ HFE formation. However, KREH2-dependent opposite control of canonical and non-canonical editing occurs in most of the ND7 3’ sequence examined in PCF. In BSF, the presence of the 3’ HFE is negligible and not modulated by KREH2. Native RESC6 complexes and total mtRNA exhibited more efficient KREH2-dependent repression of ND7 3’ than native RESC1 complexes in PCF. Importantly, negative control at the early checkpoint in ND7 3’ editing in PCF was reproduced upon *in vitro* differentiation.

### 3’ HFE formation involves the initiator gRNA-1 and a novel terminator gRNA and abolishes upstream canonical editing

To better understand how the abundant ND7 3’ HFE is created and may impact editing progression, we initially examined the top 10 most common 3’ HFE-containing sequences in RESC6-RIPs ± KREH2-RNAi. Notably, in most amplicons, all editing action had ceased upstream of the 3’ HFE (**Fig. 7**) after a short non-canonical editing junction of variable composition. Some amplicons lacked a junction entirely and were pre-edited immediately upstream of the 3’ HFE, i.e., junction length (JL) = 0. All samples without KREH2 (or KH2F1) knockdown had the same top three amplicon species, including one with JL = 0 (**Fig. 7A; Supplementary Figs. S9-S11**). Tallies of the top 100 amplicons confirmed the most common type with JL = 6 (**Fig. 7A**; the last edit site in each junction was tallied, and its percentage was given above the sequence). KREH2 (or KH2F1) knockdown generally increased the junction length in 3’ HFE-containing transcripts, and the species with JL = 0 was lost in the top ten amplicons in most samples (**Figs. 7B-C; Supplementary Figs. S9-S11**).

**Figure 7.**
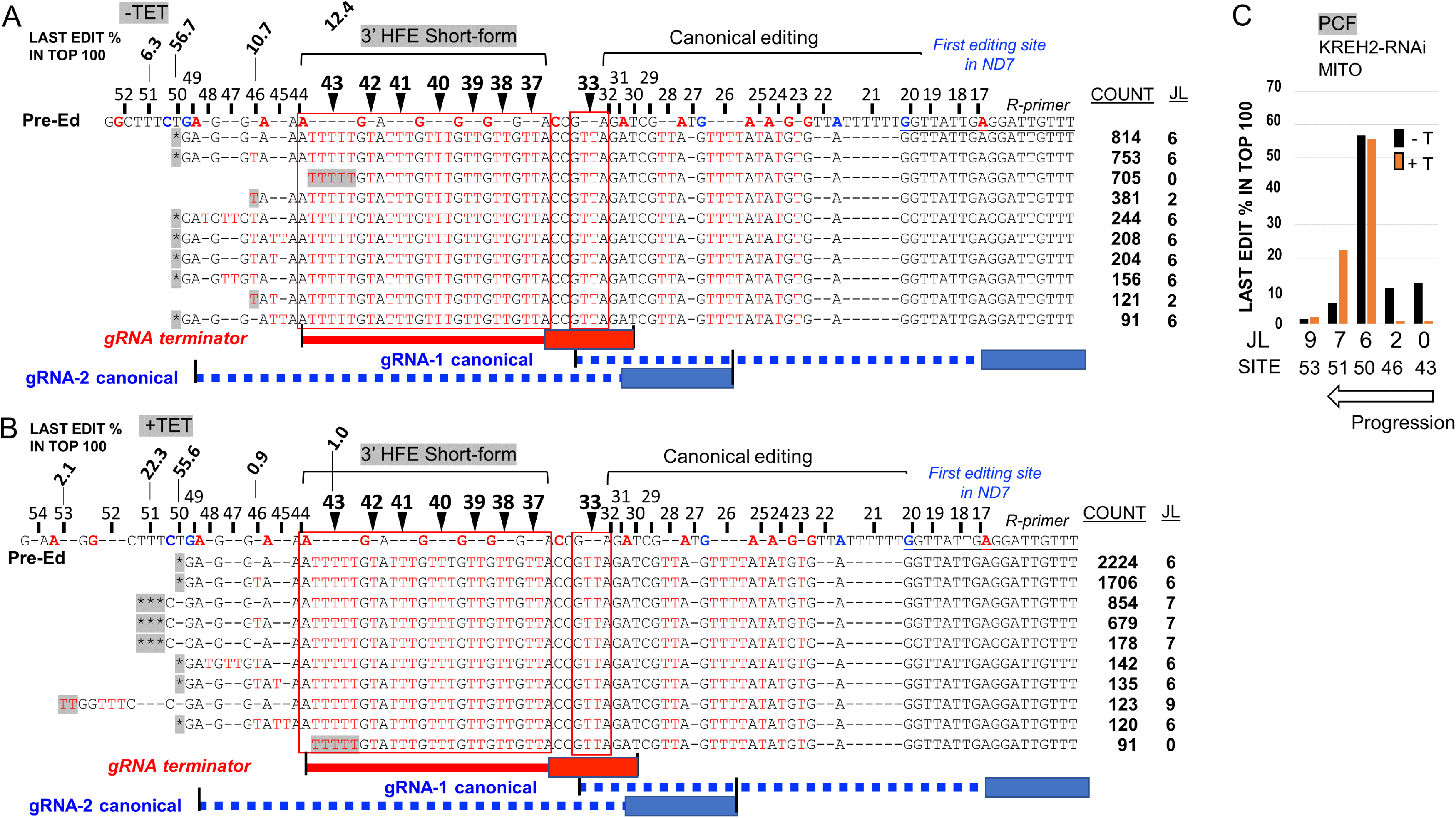
Most frequent ND7 amplicons that contain the non-canonical 3’ HFE sequence. Sequence alignment of the top ten amplicons with the short-form 3’ HFE (sites 37-43) in a representative sample of mtRNA in PCF KREH2-RNAi **(A)** uninduced -Tet or **(B)** induced +Tet. The last edit (gray) in each unique sequence is indicated as a percentage in the top 100 amplicons. Sequence 5’ to the last edit is pre-edited or includes a variable length non-canonical editing junction. The count of each amplicon in the top 100 amplicons in that sample examined is shown. The cognate initiator gRNA-1 and gRNA-2 with an anchor (box) and guiding region (dashed line) in blue, a terminator gRNA matching the 3’ HFE (straight line) in red, and the first canonical editing site in ND7 3’ (position 17) are depicted. Color-coded letters are as in prior figures. **(C)** Junction length (JL) from panels A and B, counted as the number of sites containing any edits upstream of site 43. The site containing the last edit in the junction is indicated. Amplicons carry canonically edited sequence 3’ to the 3’ HFE. Junction length (JL) analyses of other samples were also performed (**Supplementary Figs. S9-S11**).

The 3’ HFE would prevent anchoring by upstream cognate guides. Accordingly, bioinformatic searches showed that 3’ HFE-containing amplicons in our samples lacked canonical editing of blocks 2-to-4. Notably, the 3’ HFE abuts a fully edited 3’ terminal sequence in mature ND7 3’ (13,42). Combined, this 3’ terminal canonical sequence plus the 3’ HFE make a consensus extended element through site 43 (≥54-nt): 5’- AuuuuuGuAuuuGuuuGuuGuuGuuACCGuuAGAuCGuuAGuuuuAuAuGuGAG-3’. Multi-sequence alignments of the top 100 amplicons bearing the 3’ HFE confirmed the presence of the ≥54 nt extended element with minor differences. This observation suggests that this extended element derives from concerted guiding events. We noted that the 3’ HFE length suits the combined average sizes of the guiding and anchor regions in a typical gRNA (i.e., 20-to-40 nt and 6-to-11 nt long, respectively; (8,9). A search in the annotated minicircle genome from *T. brucei* BSF strain EATRO 1125 (8,9) identified a gRNA that matches the 3’ HFE (**Fig. 8A**; transcript #1). Notably, this gRNA anchor would bind, via Watson-Crick base-pairing, to a sequence created by the non-canonical +2U at site 33, while its guiding region would direct installation of the remaining 3’ HFE sequence.

**Figure 8.**
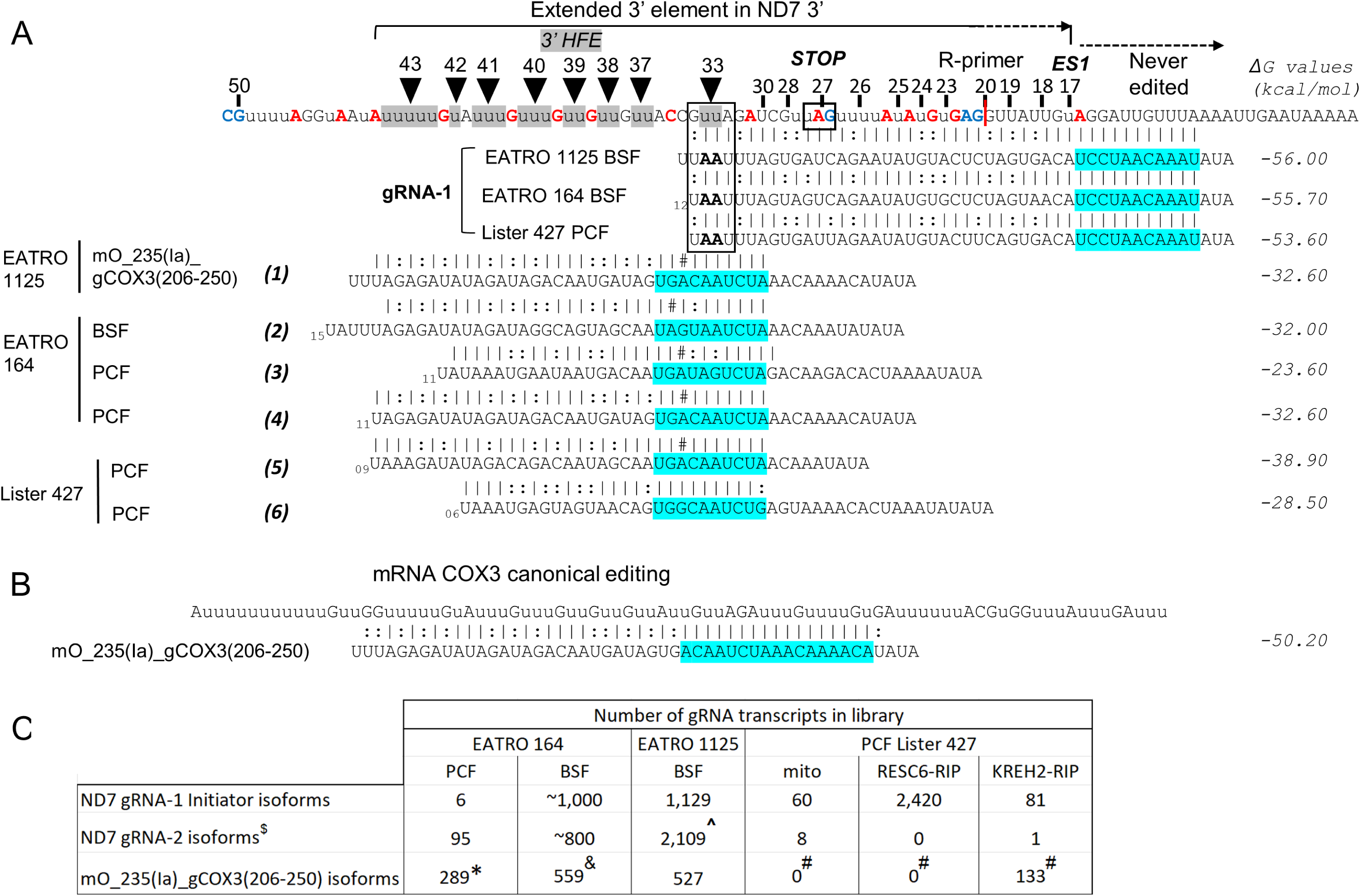
Coordinated non-canonical usage of the initiator gRNA-1 and a novel terminator guide may install the extended 3’ element in ND7 3’. **(A)** The extended 3’ element from sites 17 to 43 in ND7 3’, as in Fig. 5D, aligned with gRNA-1 isoforms in strains BSF EATRO 1125, PCF EATRO 164, and PCF Lister 427 (used in this study) with non-canonical +2U at site 33 (boxed). A novel gRNA mO_235(Ia)_gCOX3(206-250) identified in strain EATRO 1125 (transcript #1) or isoforms in other strains (transcripts #2-6) match the 3’ HFE except for a single C:A mismatch (either C in a CC doublet) at site 35 or 36 in mRNA. Guide isoform #6 in strain Lister 427 has no matches but ends prematurely at site 41 in the 3’ HFE. **(B)** Sequence alignment of gRNA mO_235(Ia)_gCOX3(206-250) with its cognate mRNA COX3. Predicted ΔG values in kcal/mol) of the duplexes are indicated. The original stop codon created by gRNA-1 in ND7 3’, a never-edited sequence, the reverse (R) primer 5’ end (red line), and anchor regions (blue) are annotated. **(C)** Number of gRNA transcripts in available databases in PCF and BSF. Initiator gRNA-1 isoforms in EATRO 164, gA6 (774-822), and Lister 427, gA6 *B1.alt*, have the same guiding capacity as in EATRO 1125. gRNAs in EATRO 164 and EATRO 1125 were determined in total mtRNA; gRNAs in Lister 427 were determined in RESC6-IPs and available alignments online (31). Transcripts no. 2 (&), no. 5 (#), and no. 4 (*). gND7(1240-1268) *B2.alt* ($), and mO_094(II)_gND7(1186-1129) (^).

Surprisingly, this gRNA is the previously classified cognate gRNA mO_235(Ia)_ gCOX3(206–250), termed hereon gCOX3, that directs editing in mRNA COX3 (**Fig. 8B**) (8,9). Similar gCOX3 guides are found in other databases in PCF strains EATRO 164 (10) (transcripts #2- to- 4) and Lister 427 (transcripts #5 and #6). Equivalent gCOX3 gRNAs were also found in PCF strains TREU 667 and TREU 927 (Donna Koslowsky, personal communication). Most of these guides (transcripts #1-to-5) have a single C:A mismatch with a cytosine dimer in mRNA. This C/A polymorphism and mismatch could potentially cause a non-canonical +1U insertion at sites 34 or 35. However, these sites exhibited relatively low NC/C values (**Fig. 4A**), suggesting that the alternative +1U edit is rare. One isoform (transcript #6) completely matches the 3’ HFE but lacks information for sites 42-43. The conservation of gRNA isoforms in multiple *T. brucei* strains and their complementarity to the extended 3’ element suggests that the proposed concerted non-canonical action by the cognate initiator gRNA-1 and non-cognate gRNA gCOX3 is biologically relevant in ND7 3’.

To assess whether gRNA relative frequency may influence the observed levels of 3’ HFE, we examined available Illumina sequenced gRNA-1 and gRNA-2 for ND7 3’ and gCOX3 in several strains (**Fig. 8C**). In PCF strain EATRO 164 mtRNA, the relative abundances of gRNA-2 and gCOX3 isoforms were similar to each other but higher than that of gRNA-1. All three gRNAs appeared to have higher abundances in this strain in BSF vs. PCF. In BSF strain EATRO 1125 mtRNA, these three gRNAs exhibited comparable frequencies. In PCF strain Lister 427 mtRNA and RESC6-RIPs, gRNA-1 was detected in all libraries but appeared relatively enriched in native RESC6 complexes. Interestingly, gCOX3 isoforms were only detected in KREH2-RIPs. In summary, the relative frequency of gRNAs did not explain the observed differences in 3’ HFE abundance between our samples.

We also asked whether the stability of gRNA/mRNA pairing may influence the observed levels of ND7 3’ HFE. Predicted ΔG values of continuous duplexes suggested that the pairs between gCOX3 isoforms and noncognate 3’ HFE-bearing mRNA (**Fig. 8A)** are thermodynamically less stable than pairs between gCOX3 (**Fig. 8B**) or gRNA-2 (**Fig. 5A**) and their cognate mRNAs. However, the efficiency of editing directed by gCOX3 is significantly higher than that directed by gRNA-2 in ND7 3’ in PCF (*P*<0.005) (**Figs. 5E-F**; compare 3’ HFE vs. block 2 edited samples without KREH2 or KH2F1 knockdown). Thus, gRNA/mRNA pairing stability did not explain the observed differences in 3’ HFE abundance between our samples. These observations, together with the PCF-specific KREH2-dependent upregulation of 3’ HFE, argue against random off-targeting of mO_235(Ia)_gCOX3(206–250) isoforms on ND7 3’ but rather suggest a fixed event in evolution.

To further examine the *in vivo* action of the proposed novel gCOX3, we looked for covalently formed bimolecular chimeras between gCOX3 and ND7 mRNA. Chimeras validate on-target gRNA-mRNA pairs *in vivo* but are not considered true editing intermediates (69–72). We readily isolated gCOX3 chimeras with mRNAs ND7 and COX3 (**Supplementary Fig. S12**). Chimeras with both mRNAs appeared to use the same gCOX3 isoform (#5) in strain Lister 427, based on their sequenced 3’ terminus. Thus, the novel gCOX3 may be bifunctional, that is, playing a dual role, canonical and non-canonical, in COX3 and ND7 3’ editing, respectively. Taken together, these data are consistent with a PCF-specific major early control checkpoint in ND7 3’ editing where KREH2 normally promotes the use of the inhibitory non-cognate gCOX3 over the cognate gRNA-2. Accordingly, KREH2 knockdown reverses this preferential guide selection. The observed differences in the ND7 3’ HFE level did not correlate with relative gRNA abundance nor with duplex stability of mRNA/gRNA pairs. KREH2 in PCF may normally promote gCOX3 use, introduce a constraint on canonical gRNA-2 function, or both.

### The 3’ HFE may form “repressive” RNA structure in ND7 3’ early editing

Targeting by mO_235(Ia)_ gCOX3(206-250, covers most sites typically modified by cognate gRNA-2 at the 3’ terminus of the ORF in mRNA ND7. However, the 3’ HFE sequence itself may not disrupt downstream binding and subsequent editing by gRNA-2 (**Fig. 7A**). Potential gRNA-2-directed “repair” editing could remove the 3’ HFE and replace it with the canonical block 2 sequence. Because of the 3’ HFE’s remarkable abundance, we considered that the 3’ HFE may form RNA secondary structure that somehow hinders gRNA-2 usage. To address this possibility, we applied dimethyl sulfate (DMS) mutational profiling with sequencing (DMS-MaPseq) to examine the structure of mRNA ND7, as in our prior study (38). This probing strategy uses DMS that specifically labels the Watson-Crick face of open and accessible adenine and cytosine bases in the RNA (9,22), and a high-fidelity processive thermostable group II intron reverse transcriptase (TGIRT) enzyme (73). We compared full-length *in vitro* T7-transcribed ND7 in pre-edited form or the most frequent 3’ HFE-containing molecule in all samples (**Fig. 7; Supplementary Figs. S9-S11**). We used the DMS reactivities as folding constraints to generate structural models of the ND7 3’ terminus (**Figs. 9A-B**). These models showed that the entire 3’ HFE and the binding site for cognate gRNA-2 in the ≥54 nt extended 3’ element form a semi-continuous helical region. This proposed stable fold may reduce access to the binding site for gRNA-2 and thereby “repair” editing by this guide. In contrast, equivalent positions in pre-mRNA appear largely accessible to DMS reactivity. DMS reactivity profiles showed particularly low signals in the predicted fold, including the 3’ HFE (sites 33-43) and binding site for the gRNA-2 anchor, supporting the hypothesis that these features are occluded in ND7 3’, whereas relatively higher DMS reactivity in upstream pre-edited and downstream edited sequence, including the binding site for the initiator gRNA-1, are generally more accessible to DMS. Therefore, our DMS-MaPseq determined structures *in vitro* support a model whereby KREH2-promoted non-canonical action by the non-cognate terminator gCOX3 and 3’ terminus of the cognate initiator gRNA-1 might form an RNA fold that sequesters the mRNA 3’ terminus. The proposed allosteric blockage of downstream cognate guiding would prevent the removal of the 3’ HFE to restore editing progression.

**Figure 9.**
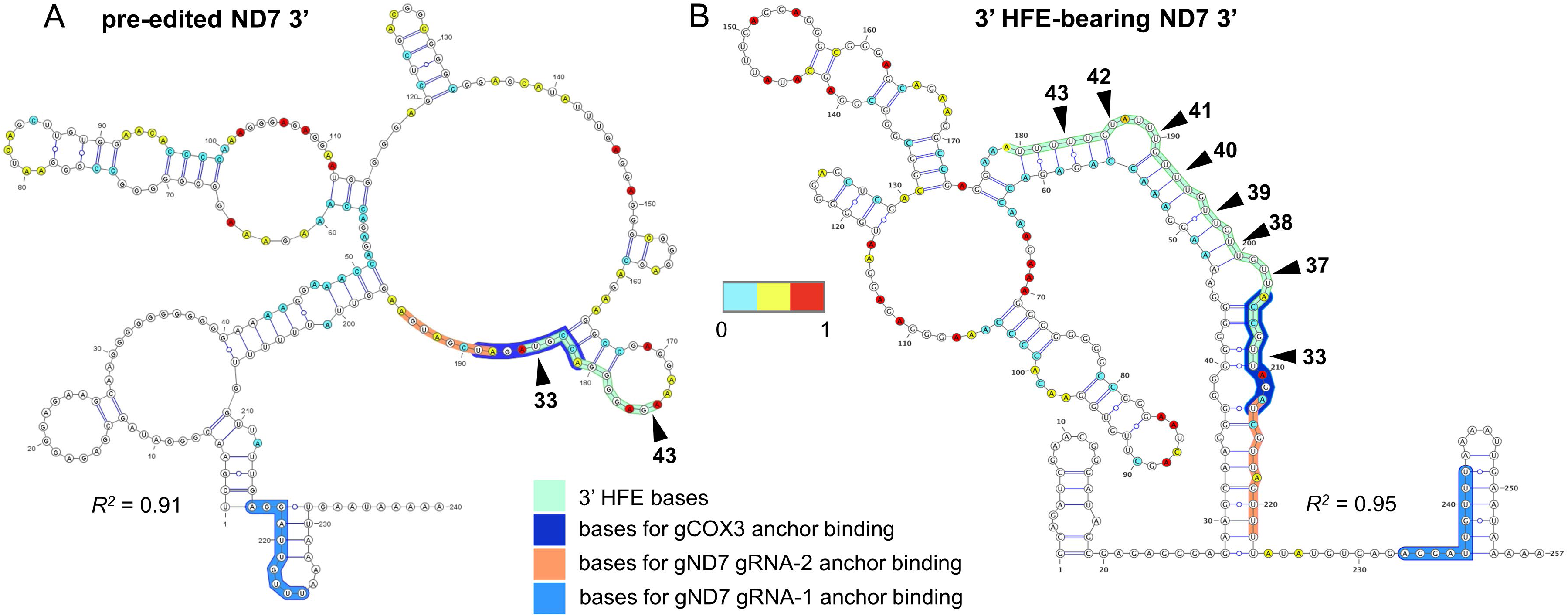
Structural models determined by DMS-MaPseq of the (A) pre-edited (PE) and (B) 3’ HFE-containing ND7 3’ terminus. Nucleotides are color-coded by normalized DMS signal. The 3’ HFE is highlighted in a green box; nucleotides used in binding the anchor region in cognate ND7 gRNA-1 and gRNA-2 and non-cognate gCOX3 are highlighted in blue, orange, and royal blue, respectively. Both structures carry a complete anchor binding site for cognate gND7 gRNA-1. Only the 3’ HFE-containing structure carries a complete element (sites 33-43) and complete anchor binding sites for cognate gND7 gRNA-2 and non-cognate gCOX3. *R^2^* is Pearson’s *R^2^* in structures from n = 2 experiments. DMS reactivity is calculated as the ratiometric DMS signal per position normalized to the highest number of reads in the displayed region, which is set to 1.0.

## DISCUSSION

Editing mediated maturation at the 3’ of the mRNA encoding ND7 is typically more efficient in BSF than in PCF cells, and is downregulated upon induced *in vitro* differentiation from PCF into mammalian infective BSF forms (14,44,46). However, the key regulatory factors driving stage-specific or preferential RNA editing control have remained a mystery. This study supports a PCF-specific repression model for ND7 3’ editing that requires KREH2 (**Figure 10**). In this model, KREH2 concurrently downregulates canonical editing and upregulates gRNA-programmed non-canonical editing in ND7 3’ in PCF cells. PCF-specific repression preferentially occurs at a major early checkpoint, where a novel “terminator” gRNA installs a 3’ HFE and a proposed repressive structure. Remarkably, KREH2 RNAi-knockdown in PCF relieved repression by reversing the two editing types thus confirming the role of KREH2 in PCF-specific ND7 3’ negative control. KREH2 is the first identified protein to repress canonical editing in any mRNA substrate during development. While KREH2 reduces the editing fidelity of ND7 3’ in PCF, KREH2 was shown to increase editing fidelity in mRNAs A6 and RPS12 in PCF (13,38). Thus, KREH2 exhibits a dual role, acting as a repressor or activator in a substrate and stage-specific manner. This role likely involves KREH2-dependent selection and utilization of cognate vs. regulatory non-cognate gRNAs, which may “tune” gene expression in either direction. Uncovering that KREH2 is not only a key protein in developmental editing control but shows a dual role as a regulator is unprecedented in *T. brucei*.

**Figure 10.**
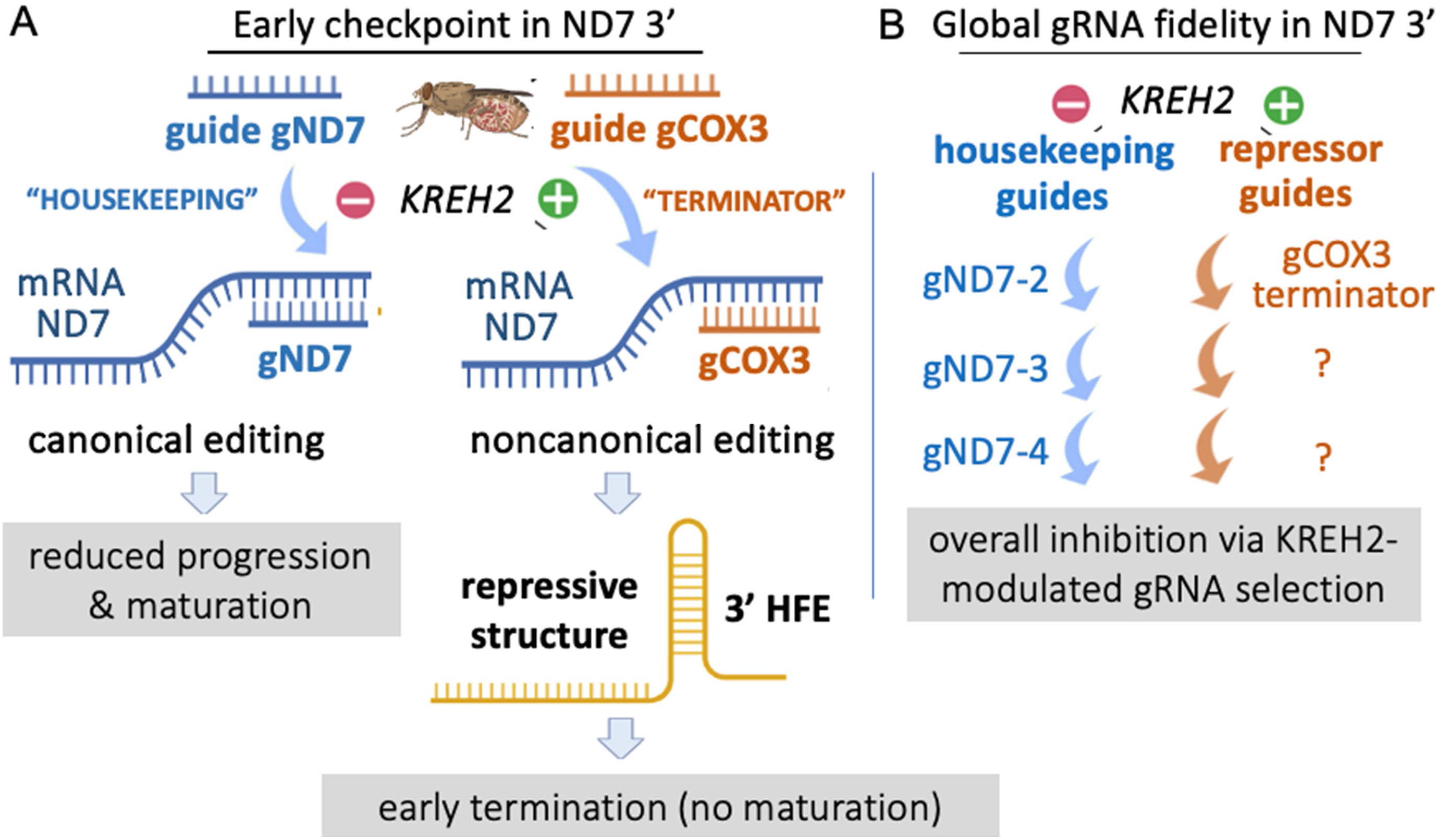
Model of REH2C-dependent developmental ND7 editing regulation via modulation of gRNA selection and RNA structure. In this model, KREH2 maintains a state of ND7 3’ editing repression in PCF involving modulation of gRNA selection and installation of a proposed RNA structure. PCF-specific repression was defined at two levels: **(A)** at a major early editing checkpoint where KREH2 exhibits concurrent positive (+) control on usage of a novel “terminator” guide (gCOX3) and negative (-) control on usage of the canonical cognate gRNA-2 (gND7). KREH2 promotes preferential usage of the terminator over gND7. Editing using the terminator installs a non-canonical 3’ HFE in ∼30% of the ND7 transcriptome, which has two main effects: first, it derails all upstream canonical editing, and second, it forms a stable fold, potentially hindering downstream gRNA-2 binding and “repair” via re-editing to replace the 3’ HFE with canonical sequence, and **(B)** globally, with gRNA fidelity changes by KREH2 promoting “repressive” non-cognate guiding and inhibiting canonical cognate guiding at most ND7 3’ sites examined. Thus, inhibition of ND7 3’ editing maturation in PCF is mainly at the early checkpoint but also occurs globally in transcripts that did not receive the 3’ HFE. KREH2-mediated repression in ND7 3’ was not observed in BSF cells.

### A PCF-specific major checkpoint in early ND7 3’ editing repression involves KREH2-dependent inhibition of cognate and promotion of non-cognate gRNA usage

The major early checkpoint in ND7 3’ editing may involve constraints on cognate guiding and or enhanced non-cognate guiding that out-competes the former. A two-step model of KREH2-promoted non-canonical editing by the initiator ND7 gRNA-1 and a novel terminator COX3 gRNA may install the 3’ HFE. In the first step, two 3’ terminal adenosines in ND7 gRNA-1 that abut the classified guiding region direct the most frequent non-canonical edit (+2U at site 33) examined in ND7 3’ (**Supplementary Fig. S6**). In native RESC6-complexes, this +2U event accounted for ∼94% of all non-canonical reads (∼60% of all reads at site 33). This +2U event led to the most robust KREH2-promoted editing pausing at site 32 (PPS32) in the ND7 3’ sequence examined in PCF. Paradoxically, strong pausing at site 32 requires highly efficient block 1 maturation, i.e., edited block 1 is generated in >70% or >75% of the PCF ND7 transcriptome’ in mtRNA or RESC6-IP samples, respectively (**Fig. 6A**). Restoring canonical block 2 editing during development in BSF cells would necessitate either reduction of the +2U edit by gRNA-1, promotion of cognate gRNA-2 function over the non-cognate terminator, or both.

Like ND7 3’, the initiator gRNA-1 in mRNA RPS12 has two conserved 3’-terminal adenines that direct a dominant non-canonical edit modulated by KREH2 (13). Thus, specific bases outside the classified canonical guiding region in some initiator gRNAs may direct strong KREH2-modulated pausing in early editing. In the second step of the concerted model, a non-cognate guide gCOX3 [mO_235(Ia)_gCOX3(206–250)] installs the remaining 3’ HFE sequence in ND7 3’. Importantly, +2U at site 33 would create a binding site for gCOX3 in ∼60% of the ND7 transcriptome. About half of these molecules (∼30% of ND7 3’ reads) contained the consensus 3’ HFE in samples without KREH2 knockdown. Thus, KREH2-promoted non-canonical guiding, involving “unexpected” edits by a cognate guide (ND7 gRNA-1) and “expected” edits by a noncognate guide (gCOX3), install the consensus 3’ HFE. Besides sites 33 and 37-43 in the 3’ HFE, sites 49 and 55 and other sites also exhibited large KREH2-promoted NC/C ratios (near 1 or >1; **Supplementary Table ST3**). Thus, substantial KREH2-dependent pausing occurs at the corresponding IPS48 and IPS54, among other sites along ND7 3’.

Analyses of all amplicons in samples confirmed the absence of canonical editing upstream of the 3’ HFE. KREH2-modulated editing by non-cognate gCOX3 derails canonical editing beyond block 1, meaning that it is a terminator. Short non-canonical junctions of variable composition and unclear sources usually followed the 3’ HFE and the junction length (JL) increased in the KREH2 knockdown (**Fig. 7**; **Supplementary Figures S9-S11)**. Interestingly, reported knockdowns of several RESC proteins increased the JL immediately upstream of canonical editing in other mRNA transcripts (35,74,75), suggesting that this may be a general feature of editing factor loss-of-function .

Overall, a PCF-specific major checkpoint in early editing of ND7 3’ involves KREH2-dependent utilization of cognate initiator gRNA-1 and a novel terminator gRNA gCOX3. Surprisingly, PCF cells exploit the highly efficient initiator ND7 gRNA-1 to install the most frequent non-canonical edit in ND7 3’, which also causes the largest pause in editing progression. The lack of this repression system in BSF facilitates efficient ND7 3’ maturation, i.e., the 3’ HFE is rare (∼0.01%), and KREH2 promotes neither 3’ HFE formation nor other non-canonical editing in most sites examined in ND7 3’ in BSF. Thus, KREH2-dependent differential gRNA selection establishes negative control at an early major checkpoint in ND7 3’ editing in PCF (**Fig. 10A**).

### KREH2-dependent gRNA selection may occur throughout the editing domain in ND7 3’

While ∼30% of the ND7 3’ transcriptome in PCF carried the repressive 3’ HFE, ∼10% and 20% of ND7 3’ transcripts in mtRNA or native RESC6 complexes, respectively, lacked the 3’ HFE and thus exhibited canonical editing beyond block 1 (**Fig. 5F**; **Fig. 6A**). Notably, KREH2 knockdown in PCF significantly increased canonical editing and decreased non-canonical editing at most sites examined in ND7 3’ amplicons that lacked the 3’ HFE. This phenotype was more robust in RESC6 complexes than in total mtRNA, suggesting active modulation by REH2C in holo-editosomes. These observations have two important implications for our proposed model of ND7 3’ editing repression in PCF. First, enhanced maturation of full editing blocks upon KREH2 knockdown indicates that KREH2 controls editing by an entire gRNA guiding domain and not just of specific bases in the guiding domain. Second, KREH2-induced editing pausing and possibly termination may affect most sites along ND7 3’. So, KREH2-dependent gRNA usage, cognate and non-cognate, may occur throughout ND7 3’, not only at the major early checkpoint (**Fig. 10B**). KREH2 may also modulate editing guided by specific gRNA bases outside the classified guiding regions in other gRNAs, as shown for conserved terminal adenines in initiator gRNAs in ND7 3’ in this study, and in RPS12 in a prior study (13). Although KREH2 may control gRNA usage or even usage of specific bases in a gRNA, mRNA turnover control may also be important during development (76). The frequency of gRNAs, cognate or non-cognate, or mRNA-gRNA duplex thermodynamic stability (predicted ΔG) may not be critical determinants in developmental ND7 3’ control. Thus, KREH2-dependent utilization of cognate and non-cognate gRNAs may repress editing throughout ND7 3’ in PCF, but mostly at an early checkpoint (**Fig. 10**).

### RNA conformation in the ND7 3’ early checkpoint may hinder potential “repair” editing in PCF

The 3’ HFE appears to abolish further editing action in ND7 3’. However, cognate gRNA-2 could potentially bind downstream of the 3’ HFE and result in editing that replaces this repressive element with canonical sequence. Although “repair” editing seems feasible, the 3’ HFE exhibits a massive accumulation in PCF. To solve this conundrum, we propose that 3’ HFE removal may be blocked by at least two colluding processes in PCF. First, as discussed above, KREH2 downregulates gRNA-2 function (**Fig. 4E**; **Fig. 6A**). Second, the 3’ HFE may form a stable RNA fold determined by *in vitro* DMS-MaPseq that blocks mRNA 3’ terminus access to gRNA-2 (**Fig. 10**). This RNA fold is an extended co-axial helix that potentially hijacks the gRNA-2 anchor binding site. We propose that generation of this RNA fold (requiring canonical block 1 editing plus 3’ HFE formation) would represses early editing action in PCF. It will be important to examine the structure of mRNA folded in native conditions as RNA secondary structure may be influenced by protein binding, as well as environmental conditions during development.

### Decreased total editing action upon KREH2 knockdown contributes to the observed reduced accumulation of mature ND7 3’ in PCF and BSF

KREH2 knockdown decreased non-canonical editing and increased canonical editing at most sites examined in ND7 3’ in PCF examined by RNAseq. However, KREH2 knockdown decreases the accumulation of fully edited mature mRNAs in both PCF and BSF, as measured by RT-qPCR (38). RT-qPCR typically quantitates full editing at a 5’ block in editing substrates. This apparent conundrum may be explained by a secondary effect of KREH2 knockdown resulting in a general decrease in total editing action (**Fig. 3**). Similarly, the knockdown of core editing proteins in RESC and RECC reduces the accumulation of fully edited transcripts in PCF and BSF (5,77). The cumulative loss in total edits (canonical and non-canonical) at most editing sites likely causes a net loss in full editing in these knockdowns. A reduction in general editing action may be partly due to disruptions in editing machinery assembly. For example, the KREH2 knockdown shifted the sedimentation of RESC proteins (43).

ND7 3’ editing repression via KREH2-dependent inhibited canonical and promoted non-canonical editing, and putative repressive structure is specific to PCF. This negative control system is not seen in the BSF lifecycle stage, where KREH2 promotes ND7 3’ maturation and non-canonical editing is minimal. Mechanisms driving a functional KREH2 switch between negative and positive control modes during development could be protein-dependent or independent. In the REH2C editing complex, KH2F1 and KH2F2 associate with KREH2 in an RNA-free manner *in vivo* or in recombinant form (**Fig. 1B**) (32,38,39), but these helicase factors may exhibit functional differences in PCF and BSF (unpublished data). Additional proteins may affect KREH2 function even if they do not bind REH2C directly, but perhaps by affecting RNA substrate or ATP binding by KREH2. Temperature could also play a role in editing control, as it directly influences RNA structure dynamics (22,78). “RNA thermometers”, perhaps aided by KREH2, may regulate editing in response to shifts in temperature and intracellular conditions. Thermometers in *T. brucei*, as in other systems, including in plants and chloroplast in green alga (78,79), could influence mRNA and or gRNA secondary structure, or RNA helicase conformation, masking or promoting productive cognate or non-cognate RNA hybrids. Interestingly, temperature reduction has been shown to promote editing efficiency of some cytochrome mRNAs. In the case of COX2, expression of the accessory protein p22 and depletion of RDK1, a kinase that inhibits BSF to PCF differentiation, was coupled to cold-induced COX2 editing (68).

### Developmental regulation of a multifunctional gRNA transcriptome including moonlighting gRNAs by the helicase editing complex REH2C

The discovery of “moonlighting” gRNAs with KREH2-dependent differential utilization in ND7 3’ and in A6 (38) suggests that at least part of the gRNA transcriptome is potentially multifunctional and regulated during development. Indeed, the novel gCOX3 and gCR4 guides, which promote editing in their cognate mRNAs, also install an abundant repressive 3’ HFE in non-cognate ND7 3’ and A6 transcriptomes, respectively (38). These and other possible KREH2-dependent regulatory gRNAs are potentially bifunctional. Additional studies are required to better understand how KREH2 inhibits or promotes canonical or moonlighting gRNA utilization in a substrate and stage-specific fashion. Regardless, it is increasingly clear that KREH2 regulates canonical and non-canonical editing on a large scale to control mRNA expression in any direction during *T. brucei* development (38).

If we consider ATP consumption via RNA ligation in the basic editing reaction alone (72,80,81), maturation of the entire ND7 3’ domain would consume ∼14 times more ATP than just installing the ≥54 nt extended element, which includes the 3’ HFE (**Fig. 5D**). Early editing termination may improve general energy consumption in PCF where ND7 3’ maturation may be dispensable. A general model invoking gRNA-programmed non-canonical editing to control canonical editing and, thus, energy efficiency would be a novel feature of *T. brucei* biology (38). In ND7 3’, massive gRNA-1-programmed editing, including non-canonical editing, may titrate factors in a ‘sponge effect,’ limiting editing-based strategies to bypass early termination. Further evidence for this model was suggested by a prior study that proposed two terminator gRNAs for editing of mRNA COX3, which may introduce a short element derailing canonical editing (46). These elements were more abundant in BSF than in PCF and often included upstream junctions of variable composition (46). Whether these COX3 mRNA sequence elements are structural or regulated by specific proteins has not been addressed. Furthermore, in the kinetoplastid, *L. pyrrohocoris*, some gRNAs were also proposed to target non-cognate mRNAs. However, in this case, the resulting non-cognate editing might alter the ORF sequence without derailing upstream editing (82).

### ND7 mRNA expression during *T. brucei* development

PCF-specific KREH2-dependent repression of ND7 3’ editing is consistent with long-established preferential editing maturation of BSF mRNAs for mitochondrially encoded subunits of complex I (cI). Prior studies reported that differentiation into transmission-competent BSF parasites is associated with up-regulation of cI (20,21). Characterization of cI subunits in BSF also showed the presence of multi-subunit complexes, but whether a complete cI is assembled in BSF remains unclear (18). Surprisingly, the deletion of two cI subunits in BSF suggested that electron transfer within cI is not essential and that cI does not contribute significantly to NADH dehydrogenase activity. However, simultaneous ablation of both cI and NDH2 did result in an exacerbated BSF growth phenotype suggesting that cI may be functional but redundant with NDH2 (18). In PCF, cI was detected but may not participate in the electron transport chain (83). Our results and model for efficient ND7 3’ canonical editing in BSF and KREH2-dependent editing repression in PCF are consistent with a role for cI in BSF but not PCF. Critically, *in vitro*-induced differentiation from BSF into PCF in two strains, Lister 427 monomorphic and TREU 927 pleomorphic, in different labs in this study confirmed that coordinated changes in 3’ HFE and canonical editing levels in ND7 3’ in parental PCF vs. BSF cell lines is a bonafide phenomenon during development. It will be important to examine additional strains for 3’ HFE and canonical editing levels to determine if our findings are universal and confirm the long-established preferential ND7 3’ editing maturation in BSF. Other authors used a quantitative RT-PCR assay to examine a fragment of edited ND7 3’ in different strains and cell lines (47). This assay suggested a higher level of edited ND7 3’ in BSF EATRO164 but lower in BSF SM compared to 427-derived PCF 29-13. The latter result in that study using the same cell lines as ours appears to contradict our results using Illumina and quantitative RT-PCR for reasons that are unclear to us. However, *in vitro* differentiation from 427-derived PCF 29-13 into mammalian infective forms increased the level of ND7 3’ (46).

Overall, this work demonstrates for the first time that KREH2 is a key regulatory factor in editing during *T. brucei* development. This study defines novel concepts important to understanding KREH2-dependent developmental editing regulation (model in **Fig. 10**): (i) KREH2 and presumably other proteins in the REH2C complex repress or promote gRNA utilization (cognate and non-cognate) in a substrate-specific manner to control mRNA expression in any direction during development, (ii) KREH2 function in ND7 3’ is the first example where a defined editing protein represses editing, including by inhibition of canonical editing, in a stage-specific manner, (iii) major early checkpoints may involve repressive RNA structures installed by specific non-canonical editing, and (iv) KREH2-regulated utilization of moonlighting gRNAs that have alternative functions in editing their cognate and noncognate mRNAs suggests that the trypanosomal gRNA transcriptome may be multifunctional. Additional studies will determine whether REH2C proteins developmentally regulate editing in other mRNAs and whether major early checkpoints are common across BSF and PCF lifecycle stages.

## Supporting information

Supplementary Figures_revised version

## DATA AVAILABILITY

RNA sequencing data are deposited in the NCBI SRA under Bioproject ID PRJNA1004327.

## SUPPLEMENTARY DATA

Supplementary Data are available at NAR online.

## ACKNOWLEDGEMENTS

We thank members of the Institute for Genome Sciences and Society at Texas A&M University for their excellent assistance with deep sequencing. The Afasizhev lab kindly provided the antiserum against RESC1/2 (GAP2/1) for RNA Immunoprecipitations. The Read lab kindly provided the antisera against RESC13 (KRGG2). Zihao Chen kindly assisted with microscopy. We are very grateful to Thavy Khem, Melanie Kuwahara, and Neida Murillo for their assistance in preparing Excel files and charts.

