## Supplementary Figures_revised version for "KREH2 helicase represses ND7 mRNA editing in procyclic-stage *Trypanosoma brucei* by opposite modulation of canonical and “moonlighting” gRNA utilization creating a proposed mRNA structure"

### Supplementary Tables

| | |
|---|---|
| Sites | Any position between two non-T nucleotides (cDNA) in the reference T-stripped sequence. Sites are numbered 3’ to 5’ in the editing direction. All edits just 5’ to G, C, or A are scored. |
| T number | The number of T nucleotides scored, between 0 and 16 (cDNA) just 5’ to a G, C, or A. |
| Canonical editing site (ES) | Any position between two non-T nucleotides (in cDNA) where the T number in the canonical pattern (mature) is expected. Other sites are not modified in mature mRNA. |
| Pre-edited (PE) sequence | Transcript sequence which has the genomic encoded T number at all sites. |
| Canonical (fully edited) sequence | Transcript sequence with the exact T number at all sites in the canonical pattern by a cognate guide. |
| Cognate guide RNA (gRNA) | A gRNA which directs a block of editing events that match the published canonical pattern. |
| Alternative non-cognate gRNA | A gRNA which directs a block of editing events that differ from the published canonical pattern. |
| Canonical (C) value | The total number of reads at each site with the expected T number in the canonical pattern. |
| Non-canonical (NC) value | The total number of reads at each site with a T number that differs from the canonical pattern. |
| Total editing value | The total number of reads at each site that contain any T number, except for the T number in the pre-edited sequence. |
| NC/C ratio (editing fidelity) | A normalized value at each position, that scores relative “editing fidelity” i.e., the overall deviation from the expected canonical pattern by gRNAs classified as canonical. Low NC/C ratio reflects high editing fidelity. High NC/C ratio reflects low editing fidelity. |
| KREH2-promoted intrinsic pause sites (PPS), aka KREH2-PPS | A KREH2-Promoted Intrinsic Pause Site (PPS) aka “KREH2-PPS”. Editing progression pauses at sites immediately downstream of high NC/C values. KREH2-RNAi knockdown significantly decreases NC/C at many sites, thereby increasing general editing fidelity and progression. |
| 3’ High-Frequency Element (3’-HFE) in ND7 3’ | An abundant noncanonical element revealed by a block of high NC/C values. The 3’ HFE in ND7 3’ is installed via coordinated noncanonical use of 3’-terminal adenosines in the gRNA-1, and a novel KREH2-modulated terminator gRNA. The 3’ HFE hinders upstream canonical editing. |
| Repressive RNA fold in ND7 3’ | A stable secondary-structure determined by DMS-MaPseq made by a ~42 nt element (including the 3’ HFE), which may occlude the 3’ terminus in ND7 3’ blocking gRNA-2 use and potentially gRNA-1-mediated “repair” which could replace the structural 3’ HFE with canonical sequence. |
| KREH2-modulated terminator gRNA | A gRNA with a proposed novel dual function modulated by KREH2: e.g., canonical in cognate mRNA CO3, and non-canonical as a putative repressor (terminator) in non-cognate mRNA ND7. This KREH2-depdendent terminator guide is potentially bifunctional. |

**Supplementary Table ST1.** Glossary of terms.

**Supplementary Table ST2. Oligonucleotide and DNA construct sequences and annotations.** Table of oligonucleotides used for Illumina studies, plasmid construction, and DMS-MaPseq experiments. Links to annotated plasmid maps in benchling.com are given when relevant.

(See excel attached)

**Supplementary Table ST3. Illumina samples and statistical analyses of KREH2-dependent differences in total editing and editing fidelity (NC/C ratio) in mtRNA and RESC complexes.** Inventory of all KREH2-RNAi library samples analyzed in this study. Replicates used to calculate averages and P-values for each figure are denoted. Calculations are separated into separate tabs depending on the conditions being compared.

(See excel attached)

**Supplementary Figures**

**Supplementary Figure S1. KREH2 localization in BSF cells with kDNA and validation of a derivative cell line lacking kDNA. (A)** Immunofluorescent microscopy of BSF *T. brucei* cells, as in **Fig. 1**, shows additional examples of our localization analysis. Cells were imaged for marker proteins in editing complexes: KREH2 (REH2C) and KREL1 (RECC). White arrows point to DAPI-stained kDNA in each cell. **(B)** Genotypic analysis of our dyskinetoplastic MGA BSF strain to verify the loss of mitochondrial genome (kDNA) after a treatment with acriflavine (left). The parental SM427 strain that was not treated with acriflavine and that still carries kDNA is shown as a control (right). The presence or absence of mitochondrial genes encoding A6, and the indicated complex I proteins was confirmed by PCR, and the nuclear-encoded gene encoding LipDH was PCR amplified as a control.

**Supplementary Figure S2. mRNA ND7 3′ mature fragment in this study and comparison of total editing events in PCF and BSF. (A, upper)** Schematic of ND7 5′ and 3′ editing domains, indicating the primers (arrows) used to amplify the ND7 3′ fragment studied. The forward primer anneals to pre-edited sequence. Boxes indicate never-edited regions. **(A, lower)** ND7 3′ amplicon sequence with canonical insertions (u) and deletions (*). This edited version (out of two proposed alternative versions in the literature; see below) is found in up to ~70% and ~14% of all reads in BSF and PCF samples, respectively, in this study. This sequence may represent the "functional" version of edited ND7 3′ in our Lister 427 cells. Sites numbered 3′ to 5′ indicate spaces between two non-U nucleotides. Vertical red bars at sites 21 and 117 indicate 3′ ends of the primers used for library construction. The stop codon and 3′-UTR, including the first editing site (site 17 shaded) and never-edited region, and major KREH2-Promoted Pause Sites (PPSxx, where xx indicates the site number), are all shown. Cognate gRNAs (horizontal black bars include the first two guides that best matched our Illumina and Sanger sequenced editing data: the initiator gRNA1 ***B1*** and gRNA2 ***B2.alt*** in PCF strain Lister 427 (*29*), which are functionally identical isoforms of gND7(1269-1320) and gND7(1240-1268), respectively, in PCF EATRO 164 (*20*). (B) Alignment of representative annotated gRNAs from BSF EATRO 11125 kDNA minicircles, predicting a slight polymorphism at the C-terminus of the ORF ending in VDR (*9*). This version of ND7 3′ was found up to ~1% of all reads in our Lister 427 samples (vs. up to 70% matching a predicted ORF ending in EYR, as indicated above). (C) Site-by-site total edits in the entire ND7 3′ fragment (through site 117) in PCF vs. BSF total mtRNA (Mito, minus knockdown). Blue bars denote the first few gRNAs. Sites 33 and 37-43 of particular interest in ND7 3′ early editing are highlighted. Significant differences at each site with the average and standard deviation bars of biological replicates and *P*-values *<0.05, **<0.005, ***<0.0005. See Materials and Methods and **Supplementary Table S3** for additional details on statistical analysis.

**Supplementary Figure S3. Total editing values of ND7 3′ in mtRNA and RESC6-RIPs following PCF KREH2-RNAi and KH2F1-RNAi knockdown, and BSF KREH2-RNAi knockdown.** Site-by-site total edits in the entire amplicon in the indicated cell RNAi lines ± Tet and samples: **(A)** PCF mtRNA, **(B)** BSF mtRNA, **(C-D)** PCF RESC6-RIPs. Sites 33 and 37-43, discussed in the text, are highlighted. Blue bars denote the first few gRNAs. Other annotations and statistics are as in **Supplementary Fig. S2**.

**Supplementary Figure S4. Editing fidelity (NC/C ratio) values of ND7 3′ in mtRNA and RESC6-RIPs following PCF KREH2-RNAi and KH2F1-RNAi knockdown, and BSF KREH2-RNAi knockdown.** Site-by-site editing fidelity (NC/C ratio) in the entire amplicon in the indicated cell RNAi lines ± Tet and samples: **(A)** PCF mtRNA, **(B)** BSF mtRNA, **(C-D)** PCF RESC6-RIPs, and **(E)** PCF vs. BSF mtRNA. The highest NC/C in the amplicon at site 33 (arrowhead) caused the largest KREH2-promoted pause at site 32 (PPS32) in the sequence examined in all samples. Other annotations and statistics are as in **Supplementary Fig. S2**.

**Supplementary Figure S5. Editing fidelity (NC/C ratio) values of ND7 3′ in mtRNA and RESC6-RIPs following PCF KREH2-RNAi and KH2F1-RNAi knockdown, and BSF KREH2-RNAi knockdown.** Cumulative editing fidelity (NC/C ratio) in the entire amplicon in the indicated cell RNAi lines ± TET and samples: **(A)** PCF mtRNA, **(B-C)** PCF RESC6-RIPs. Low fidelity (high NC/C). High fidelity (low NC/C). Other annotations and statistics are as in **Supplementary Fig. S2**.

**Supplementary Figure S6. Non-canonical editing events in PCF mtRNA and RESC6-RIPs, and BSF mtRNA following KREH2-RNAi knockdown.** Site-by-site **(A-C)** and cumulative **(D-F)** non-canonical editing in the entire amplicon in mtRNA or RESC6-RIPs in PCF and BSF KREH2-RNAi cell lines ± Tet. Other annotations and statistics are as in **Supplementary Fig. S2**.

**Supplementary Figure S7. Canonical editing events in PCF mtRNA and RESC6-RIPs, and BSF mtRNA following KREH2-RNAi knockdown.** Site-by-site **(A-C)** and cumulative **(D-F)** canonical editing in the entire amplicon in mtRNA or RESC6-RIPs in PCF and BSF KREH2-RNAi cell lines ± Tet. Other annotations and statistics are as in **Supplementary Fig. S2**.

**Supplementary Figure S8. *In vitro* differentiation of BSF into PCF**. **(A)** Biomark HD Fluidigm assays of a broad range of transcripts, including hallmark mRNAs examined in **Fig. 6B,** including mitochondrial edited and never-edited, or nuclearly encoded mRNAs. The experiment was carried out in parallel using RNA extracted from monomorphic Lister 427, and pleomorphic TREU 927 cells. Undifferentiated BSF cells (d0 control) were first incubated or not with cAMP (8-(4-chlorophenylthio)-cAMP; cAMP) and grown at 37°C for two days, followed by transfer to PSF media containing CCA (citrate/cis-aconitate) and 27°C for four days (i.e., six days total following cAMP induction; d6 -/+cAMP). Formation of non-replicating BSF forms was enhanced or not, in the presence or absence of cAMP. Note that the primers targeting VSG were designed to recognize VSG221, which is expressed in the 427, but is not detected in the 927-cell line, which must therefore be expressing a different VSG. **(B)** qPCR analyses of ND7 3′ HFE, canonically edited, and pre-edited ND7 normalized to TERT as a reference transcript following *in vitro* differentiation of the pleomorphic TREU 927 cell line. Ratios in post/prior differentiation (±cAMP +CCA)/BSF are compared as for the independent experiments in Lister 427 cells that are shown in **Fig. 6D**. **(C)** Validation of ND7-HFE amplicons in qPCR assays. Sequence alignment shows the 3′ HFE (formed completely or partially) in several molecules that were amplified, cloned, and Sanger-sequenced verified. The position of the primers used in the assays is indicated.

**Supplementary Figure S9. Sequence alignment of the top ten amplicons that contain the 3′ HFE short-version in mtRNA samples from the PCF KH2F1-RNAi cell line**. **(A-B)** Sequence alignment of the top ten amplicons with the short-form 3′ HFE (sites 37-43) in representative samples of mtRNA from PCF cells where KH2F1-RNAi **(A)** was uninduced -Tet or **(B)** induced +Tet. The last edit (gray) in each unique sequence is indicated as a percentage in the top 100 amplicons. Sequence 5′ to the last edit is pre-edited or includes a variable length non-canonical editing junction. The count of each amplicon in the top 100 amplicons in that sample examined is shown. The cognate initiator gRNA-1 and gRNA-2 with an anchor (box) and guiding region (dashed line) in blue, a terminator gRNA matching the 3′ HFE (straight line) in red, and the first canonical editing site in ND7 3′ (position 17) are depicted. Color-coded letters are as in prior figures. The sequence downstream of the 3′ HFE is canonically edited. **(C)** Junction length (JL) from panels A and B, counted as the number of sites containing any edits upstream of the 3′ HFE site 43. The site containing the last edit in the junction is shaded (gray).

**Supplementary Figure S10. Sequence alignment of the top ten amplicons that contain the 3′ HFE short-version in RESC6-RIPs from the PCF KREH2-RNAi cell line. (A-B)** Sequence alignment of the top ten amplicons with the short-form 3′ HFE (sites 37-43) in representative samples of RESC6-RIPs from PCF cells where KREH2-RNAi **(A)** was uninduced -Tet or **(B)** induced +Tet. **(C)** Junction length (JL) from panels A and B. All annotations and details are as above.

**Supplementary Figure S11. Sequence alignment of the top ten amplicons that contain the 3′ HFE short-version in RESC6-RIPs from the PCF KH2F1-RNAi cell line**. **(A-B)** Sequence alignment of the top ten amplicons with the short-form 3′ HFE (sites 37-43) in representative samples of RESC6-RIPs from PCF cells where KH2F1-RNAi **(A)** was uninduced -Tet or **(B)** induced +Tet. **(C)** Junction length (JL) from panels A and B. All annotations and details are as above.

**Supplementary Figure S12. *In vivo* chimera formation of a proposed bifunctional gRNA gCOX3 with non-cognate mRNA ND7 and cognate mRNA COX3.** Multi-sequence alignment of RT-PCR amplified chimeras between gRNA gCOX3 and **(A)** HFE-bearing ND7 3′ or **(B)** canonically edited mRNA CR4. The top sequence is a reference of the predicted chimeras using the gCOX 3′ terminus of isoform #5; PCF Lister 427 in **Fig. 8**. gRNA in blue with identical 3′ bases plus U-tail captured by Sanger sequencing in both chimeras (dotted box). Common U-insertions in 3′ HFE-ND7 and COX3 mRNAs (gray shade). Forward (F) and reverse (R) primers. A drawing of chimera formation *in vivo* indicates mRNA cleavage and subsequent ligation of the mRNA 3′ fragment with the 3′ end of the hybridized gRNA.

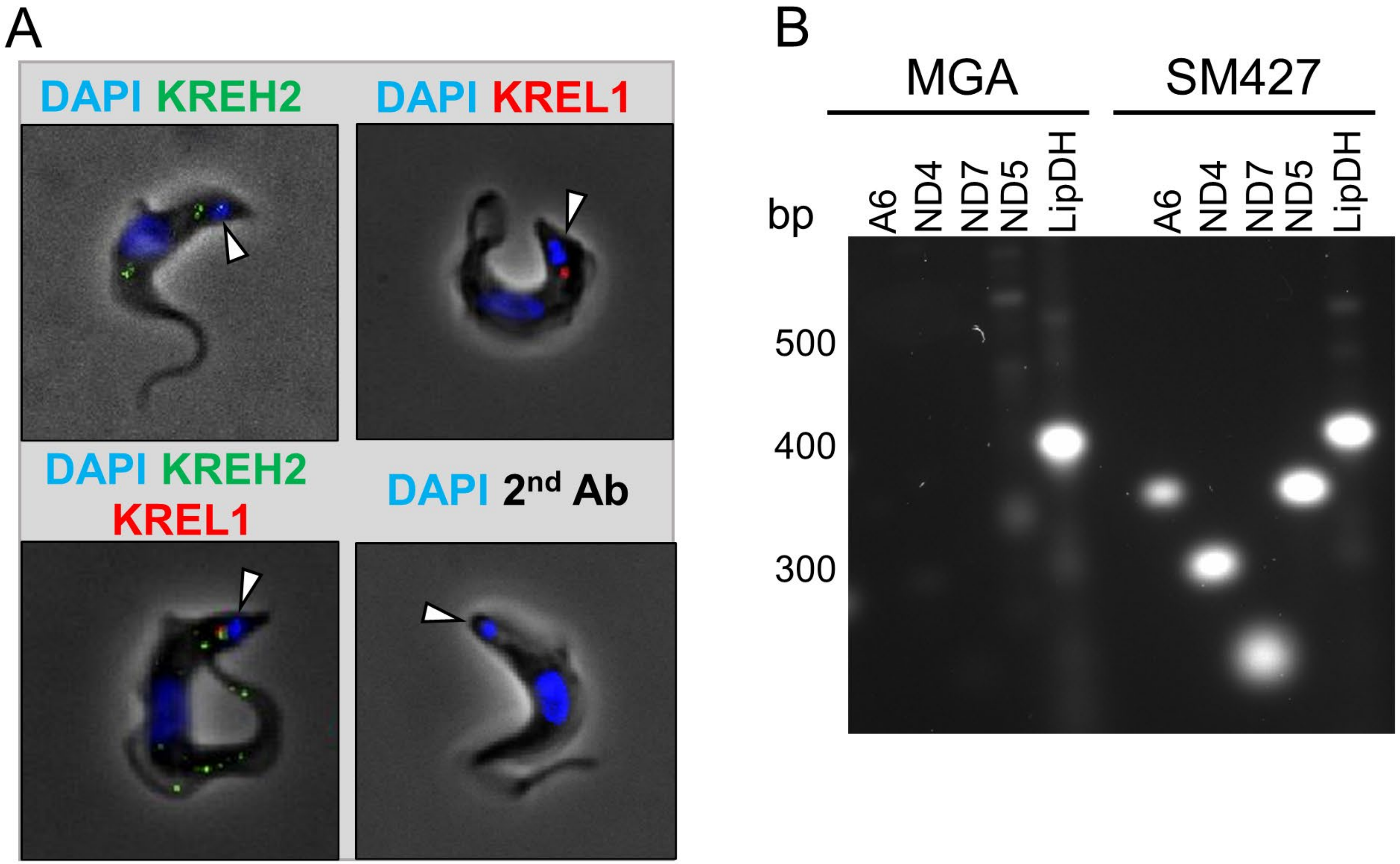


Supplemental Figure 1

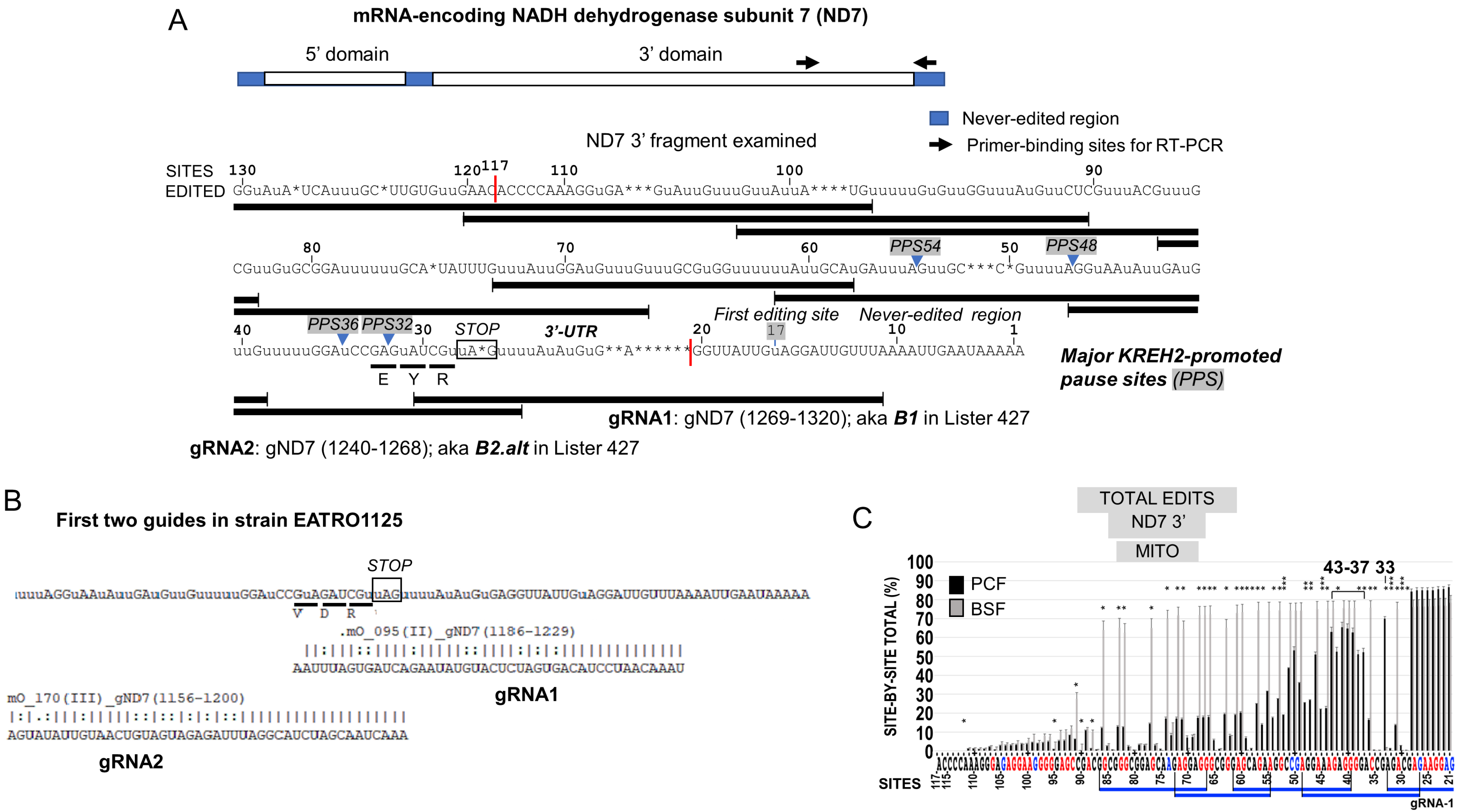


Supplemental Figure 2

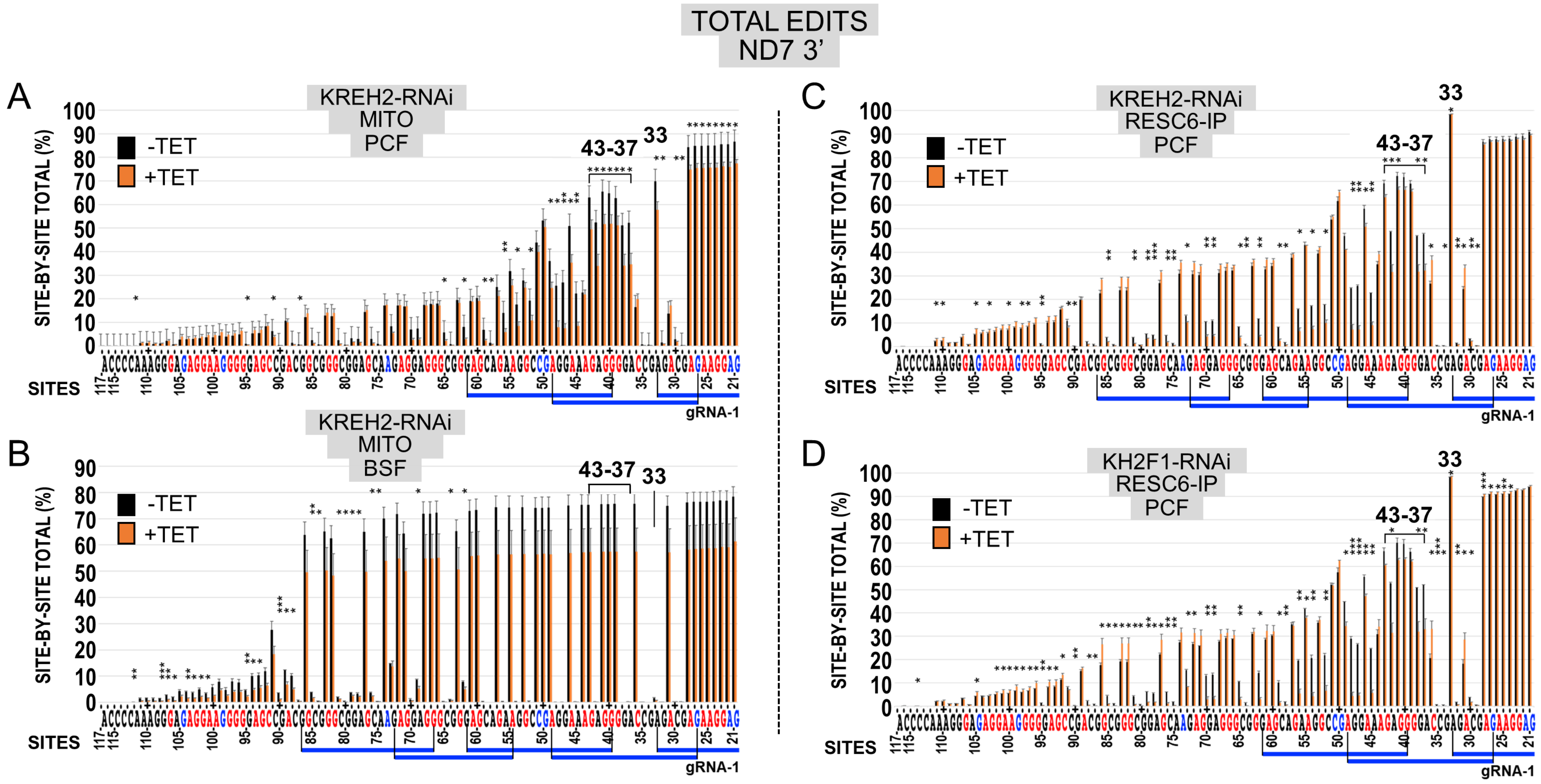


Supplemental Figure 3

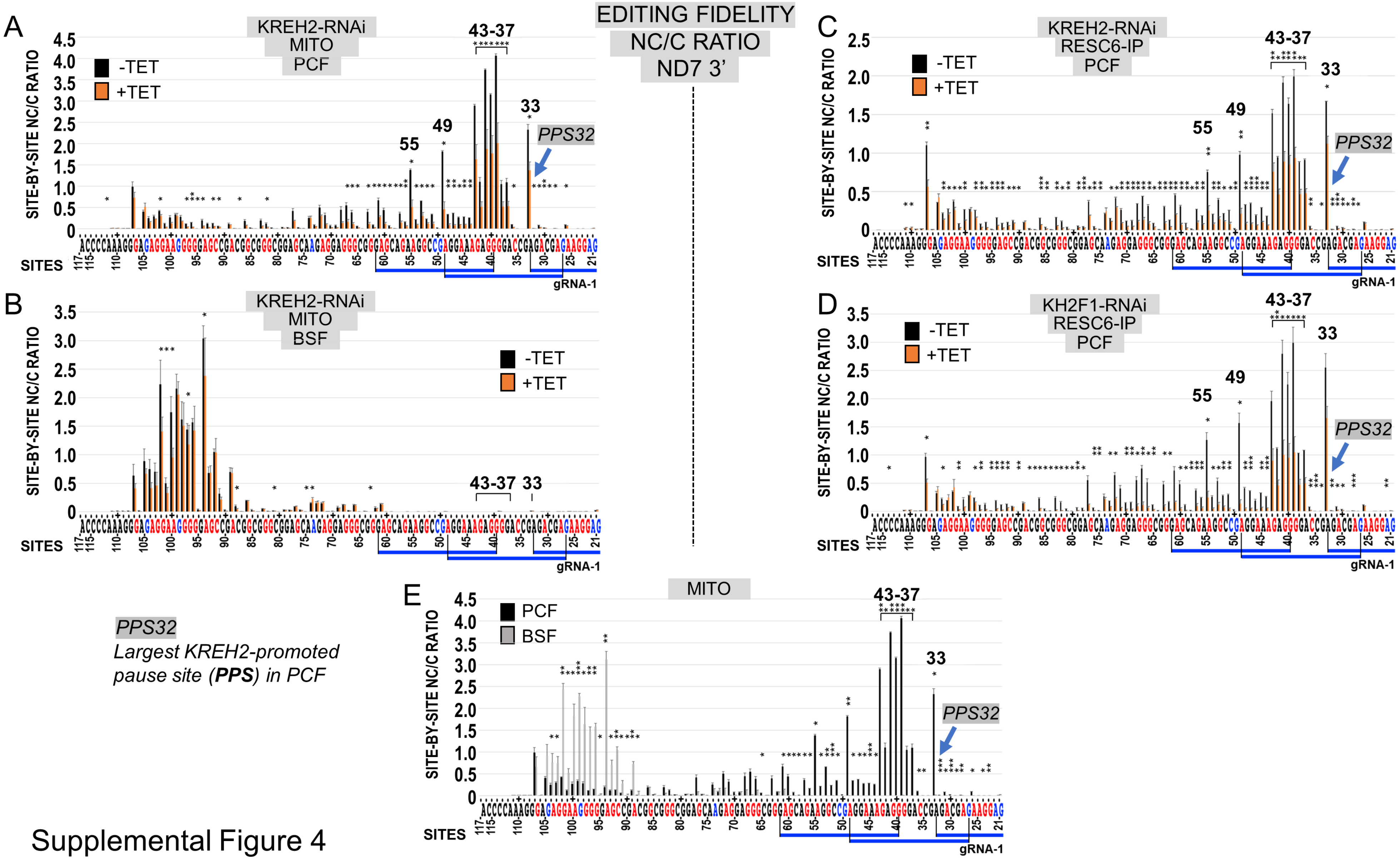


Supplemental Figure 4

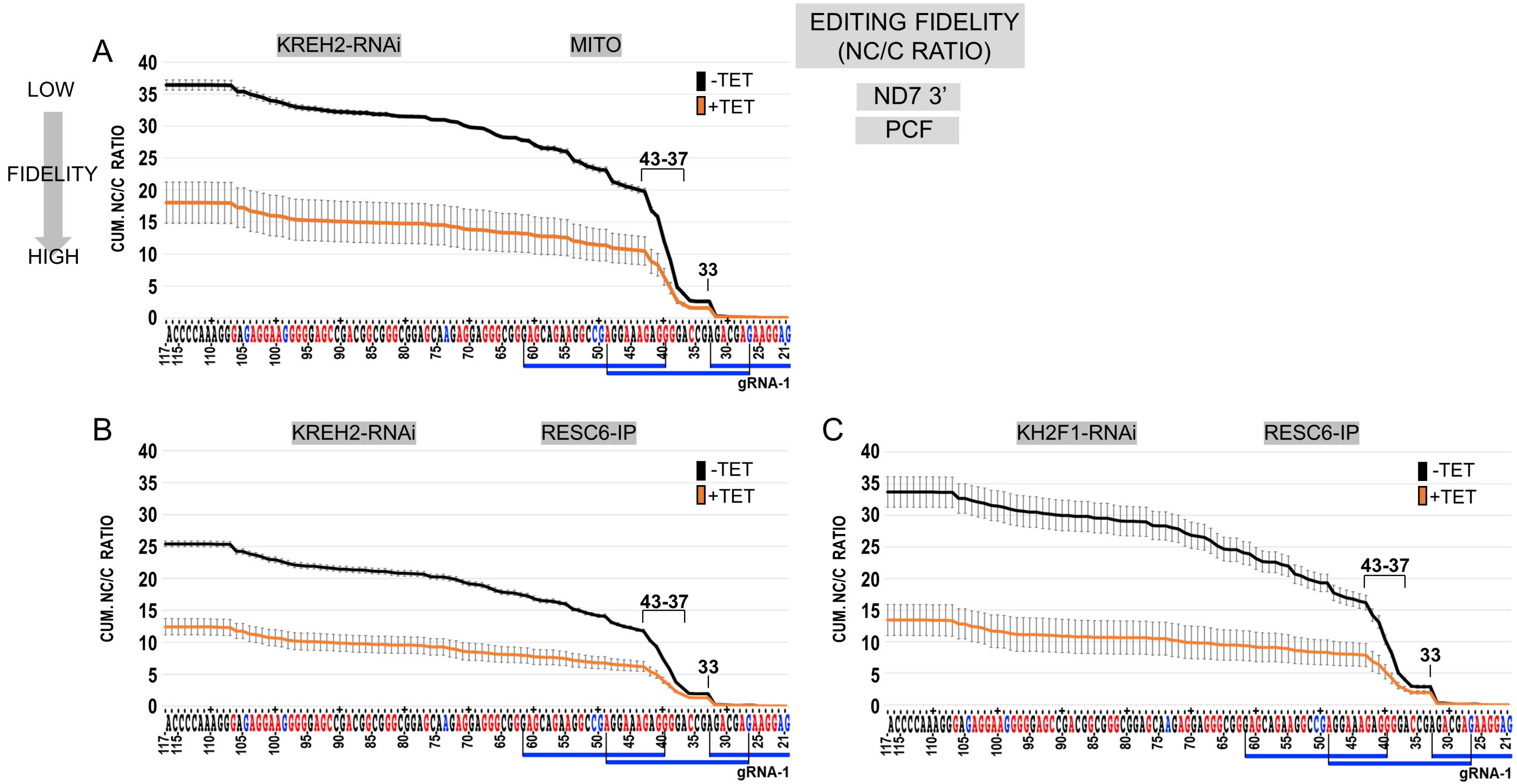


Supplemental Figure 5

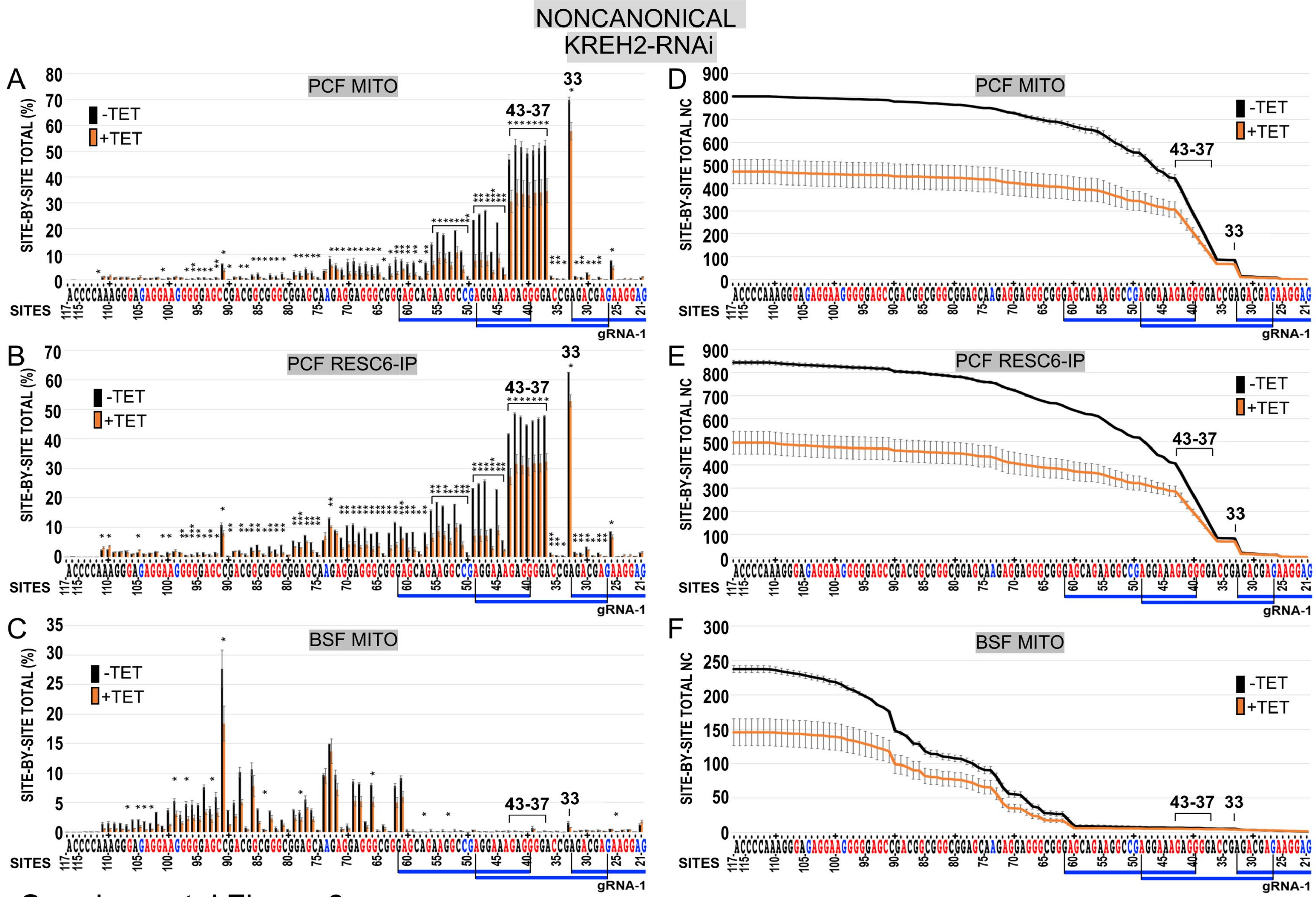


Supplemental Figure 6

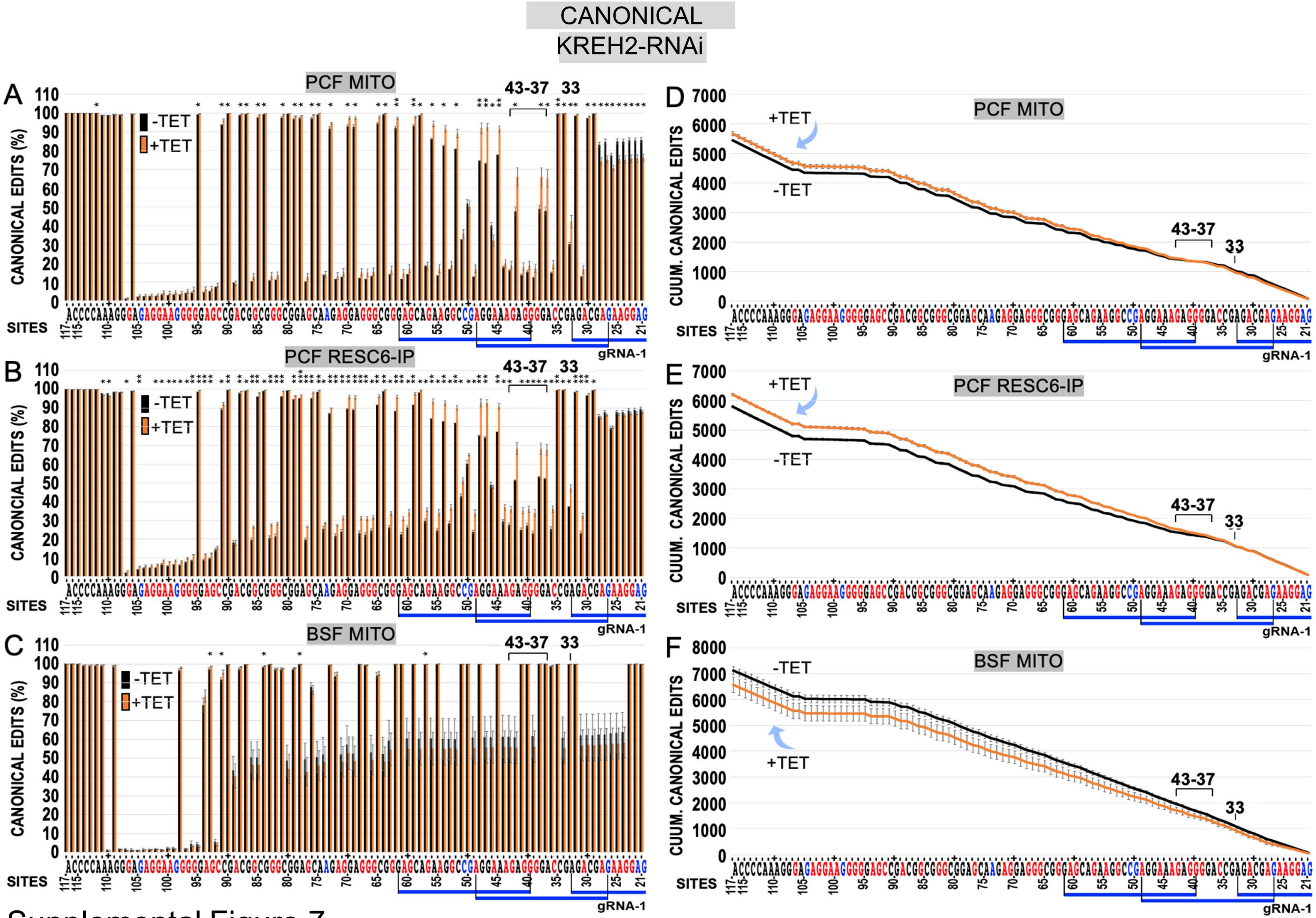


Supplemental Figure 7

*In vitro* differentiation BSF ⇒ PCF

A Biomark Fluidigm assays

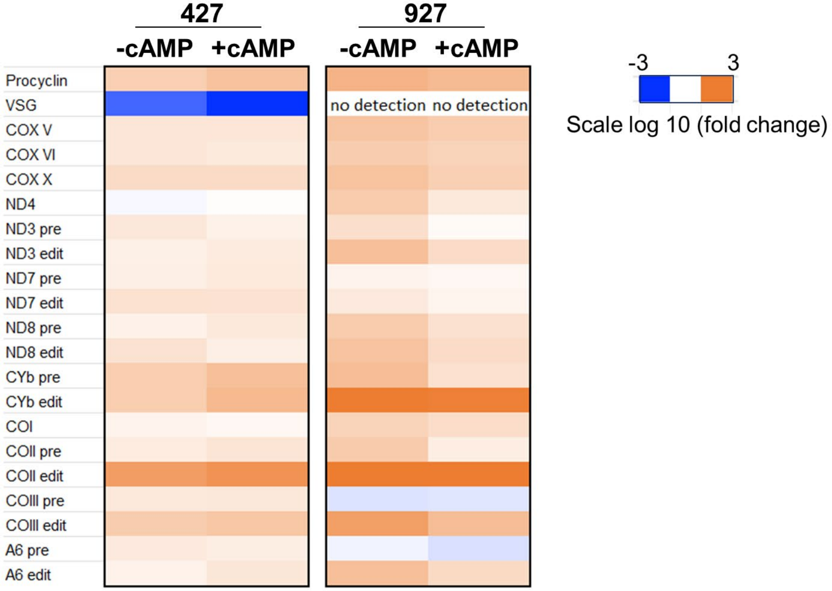


B 927

| | (+cAMP+CCA)/BSF | (-cAMP+CCA)/BSF |
|---|---|---|
| HFE | 13.96 | 75.74 |
| ED | 0.41 | 0.29 |
| PE | 1.45 | 1.93 |

| | BSF | (+cAMP+CCA) | (-cAMP+CCA) |
|---|---|---|---|
| HFE/PE | 1 | 9.61 | 39.32 |
| ED/PE | 1 | 0.28 | 0.15 |

C amplicons of ND7 with 3’ HFE

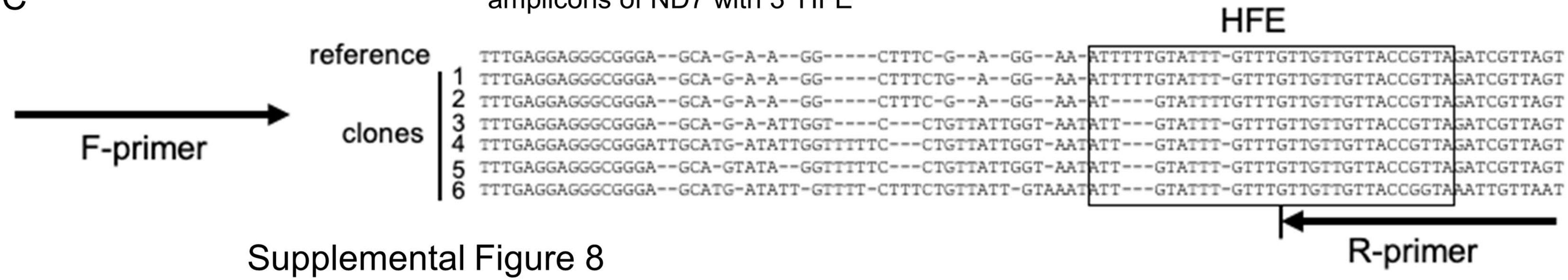


Supplemental Figure 8

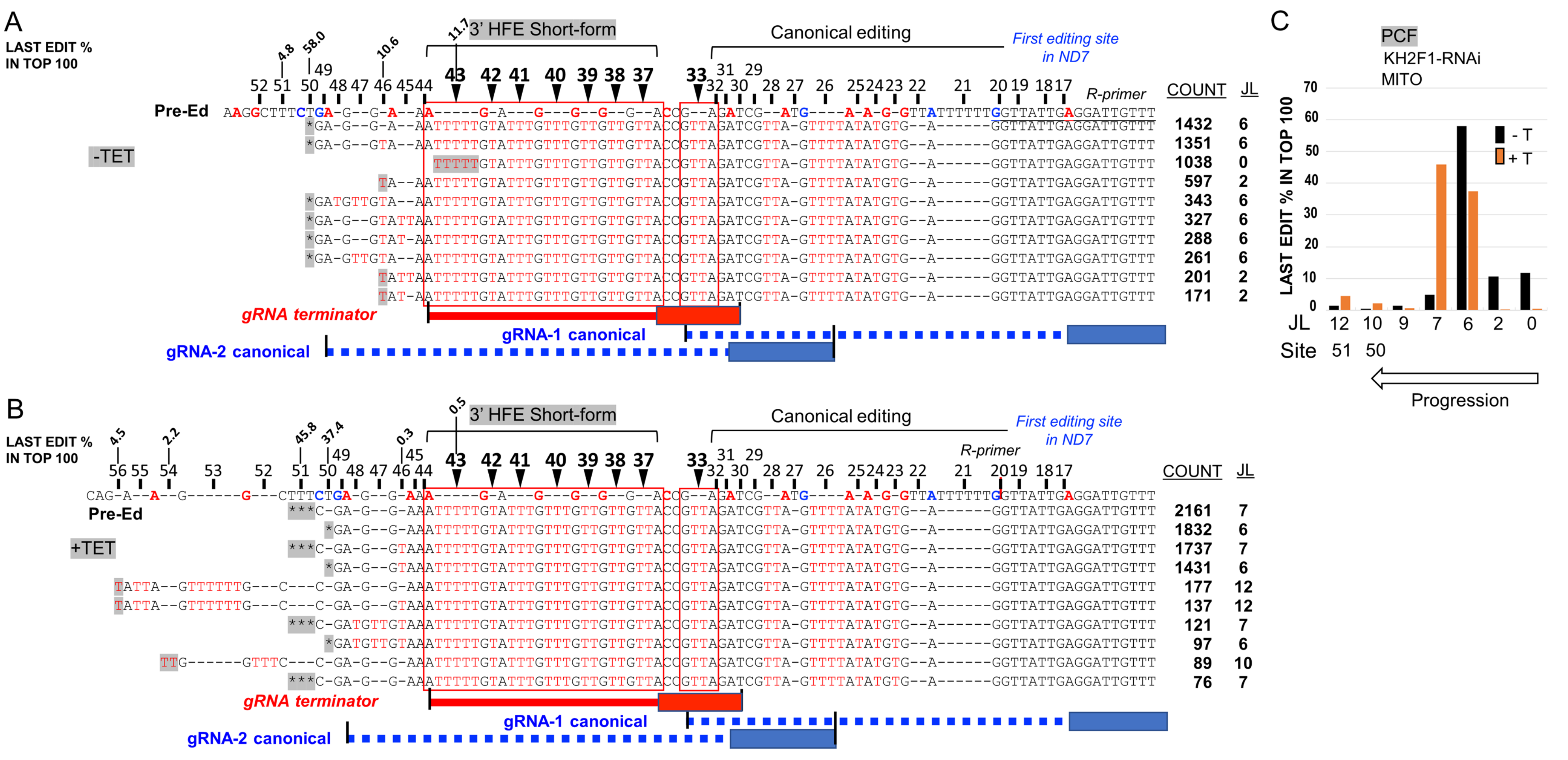


Supplemental Figure 9

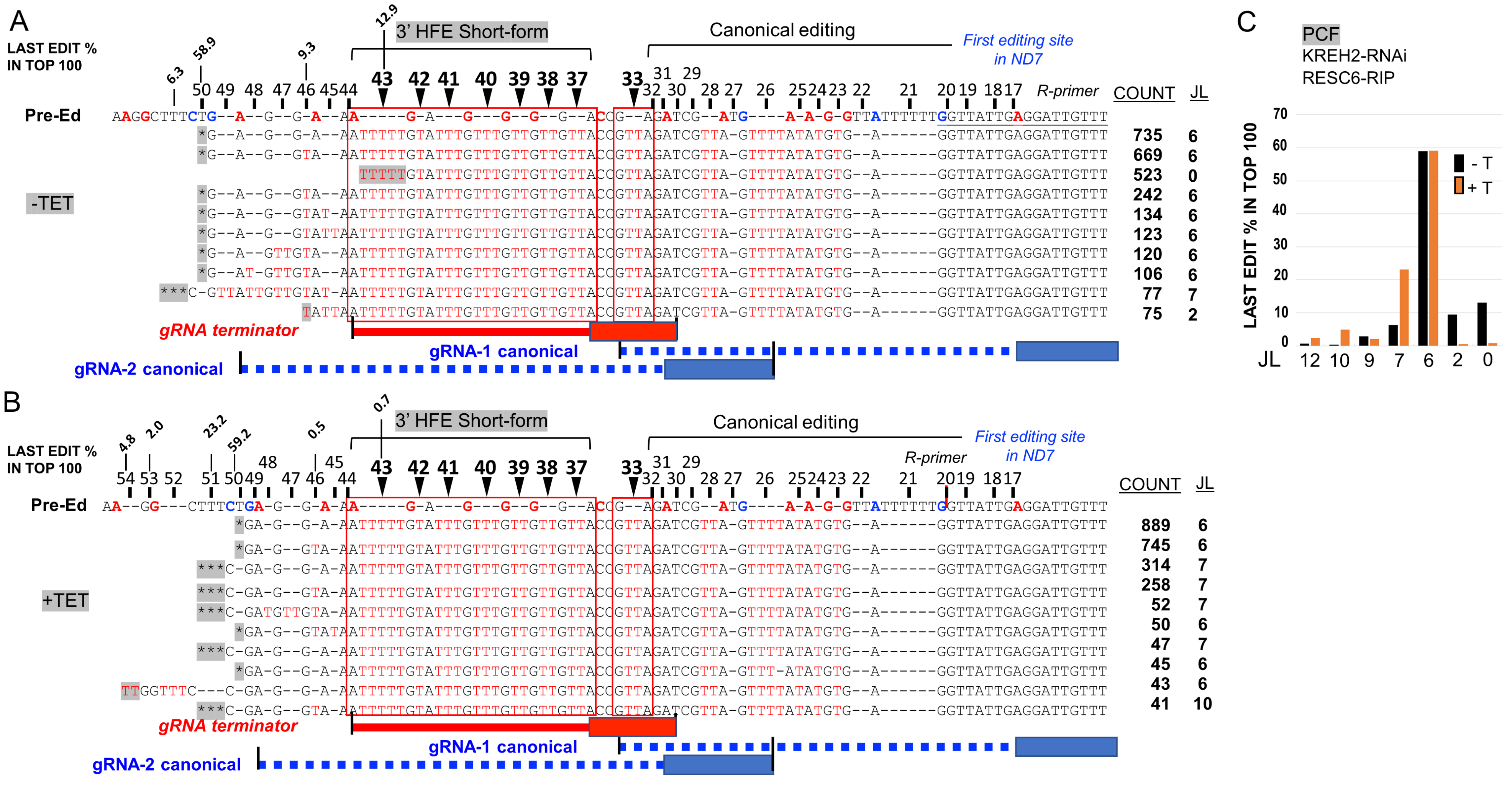


Supplemental Figure 10

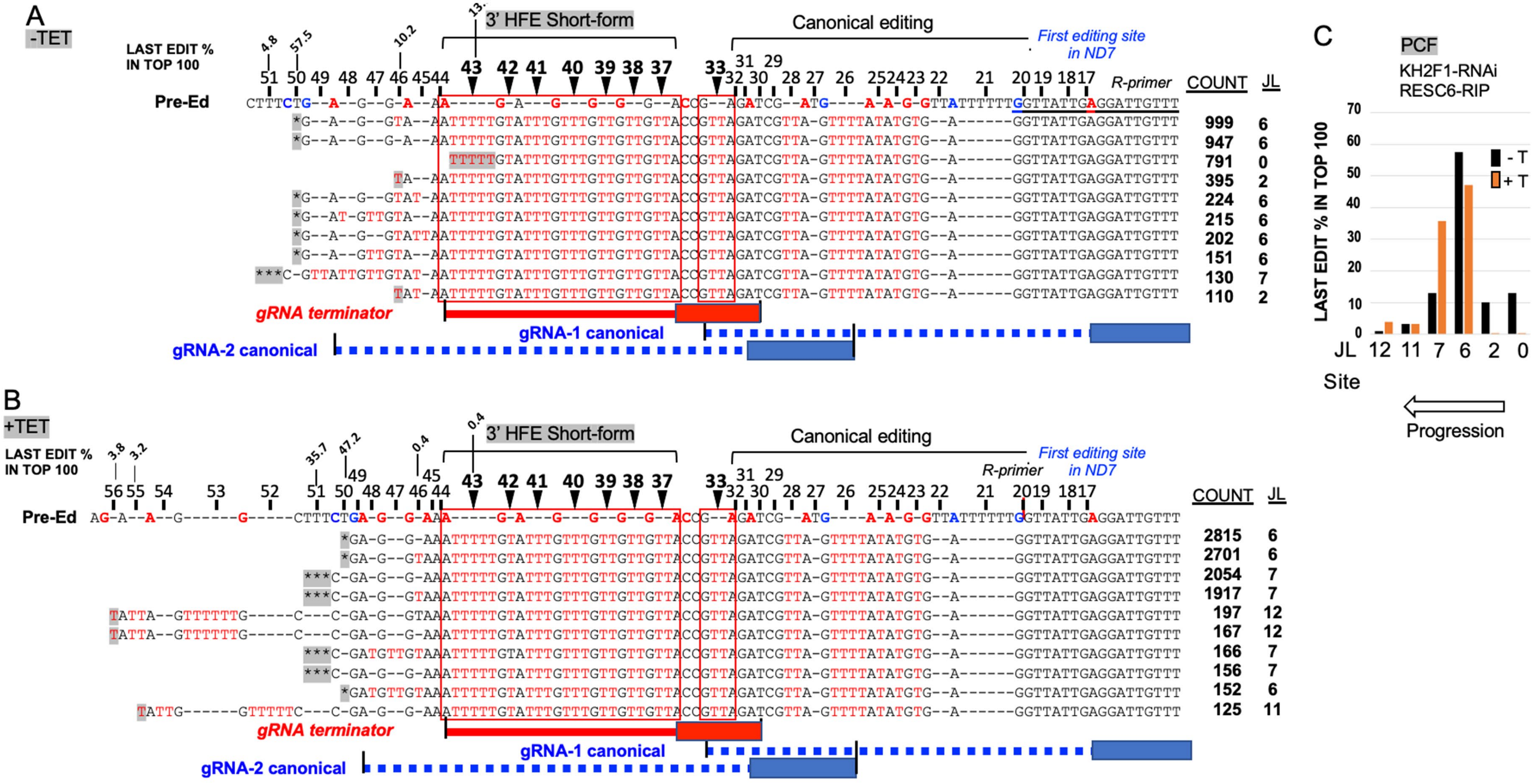


Supplemental Figure 11

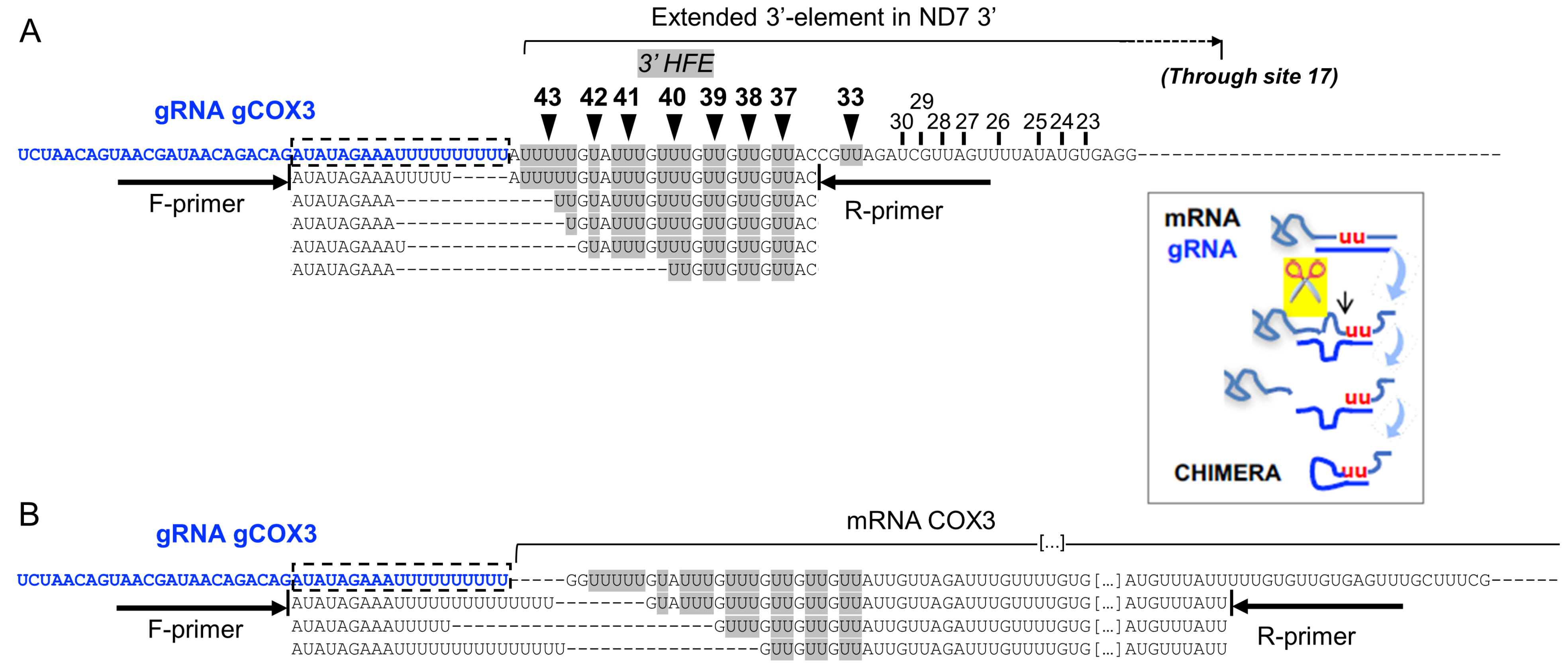


Supplemental Figure 12
